# Systematic profiling of WD40 proteins reveals Wcp1, a cyclophilin linking CO_2_/heat tolerance to acidic pH adaptation in *Cryptococcus neoformans*

**DOI:** 10.64898/2026.05.15.725558

**Authors:** Jin-Tae Choi, Seong-Ryong Yu, Jeeseok Oh, Yu-Byeong Jang, Yujin Lee, Hyunjin Cha, Doyeon Won, Dohee Kim, Sungmin Yu, Soojin Yu, Eui-Seong Kim, Seun Kang, Chaewon Kim, Kyung-Ah Lee, Jong-Seung Lee, Jaeyoung Choi, Won-Jae Lee, Kyung-Tae Lee, Yong-Sun Bahn

## Abstract

WD40 domains are major protein–protein interaction (PPI) scaffolds, yet their contributions to fungal pathogenicity remain poorly defined. We systematically analysed 94 canonical WD40 proteins in *Cryptococcus neoformans*. Conditional knockdown and sporulation identified 36 essential WD40 proteins, while in vitro and in vivo profiling of 103 signature-tagged deletion strains spanning 52 genes uncovered 31 pathogenicity-related WD40 proteins, including epigenetic and post-transcriptional regulators. We identified Wcp1, a dual-domain protein whose WD40-repeat and cyclophilin domains are required for growth at 37°C under 5% CO_2_. Its WD40 scaffold and PPIase domain supported CO_2_/heat tolerance and virulence. Notably, Wcp1 couples these functions to acidic pH adaptation: *wcp1*Δ failed to grow under elevated temperature and CO_2_ at acidic pH, exhibited enhanced intracellular acidification, reduced macrophage survival and attenuated virulence in *Drosophila* and mice. Integrated transcriptomic and proteomic analyses place Wcp1 at the centre of intracellular pH homeostasis, coordinating proton transport, metabolic adaptation and stress-buffering networks.

## Introduction

Proteins support diverse biological processes—maintaining structural integrity, catalysing chemical reactions, mediating signal transduction and coordinating immune defence—but most function not in isolation. More than 80% of proteins are thought to act as part of complexes through interactions with other proteins or nucleic acids^1–3^, underscoring the centrality of PPIs in biology^4^. Deciphering PPIs, particularly in disease contexts, is critical for therapeutic innovation. Several PPI modulators are already in clinical use or advanced trials, such as ABT-199 (targeting Bcl-2/Bax), Idasanutlin (MDM2/p53) and RVX-208 (Bromodomain/histone)^5^. A shared feature of these agents is the presence of a defined binding pocket within the PPI interface, which enables high-affinity interactions with small molecules^6–8^. By contrast, most PPI interfaces are flat and lack such druggable features, posing a major barrier to therapeutic targeting^6–8^.

Among PPI scaffolds, WD40 domains are particularly abundant in eukaryotes, ranking first in *Saccharomyces cerevisiae*, third in *Drosophila melanogaster*, and fourth in humans^9^. WD40 proteins are typically composed of seven repeats that fold into an asymmetrical β-propeller structure. Each repeat spans 44–66 residues and terminates with a conserved tryptophan-aspartate (WD) dipeptide^10^. This β-propeller architecture forms a central cavity that functions as a scaffold for PPIs. Although WD40 proteins generally lack intrinsic catalytic activity, their central cavity mediates interactions with partner proteins and may provide a favourable pocket for high-affinity binding of drug-like small molecules^11^. Despite this potential, WD40 proteins remain largely unexplored in the context of antifungal drug discovery.

A setting in which WD40-mediated PPI networks may be particularly critical is host adaptation, where pathogens must rapidly remodel their proteome to survive hostile microenvironments. Host colonisation exposes microbes to physicochemical pressures that differ sharply from external environments, including elevated temperature and host-level CO_2_. For fungal pathogens, these pressures define major barriers to disease: thermotolerance enables mammalian infection, CO_2_ tolerance has been genetically linked to cryptococcal virulence^12–15^. Yet how fungal cells integrate and buffer these co-occurring stresses at the cellular level remains poorly defined, and whether WD40-based PPI scaffolds contribute to such adaptation is unknown.

Here, we performed a systematic functional analysis of canonical WD40 proteins in *C. neoformans*, a major human fungal pathogen that causes life-threatening meningoencephalitis, with an estimated 194,000 cases with 147,000 deaths annually worldwide^16^. This survey identified multiple WD40-associated complexes implicated in cryptococcal pathogenicity, revealing a novel WD40-containing cyclophilin, Wcp1. Notably, Wcp1 promotes thermotolerance and CO_2_ tolerance by mediating intracellular pH homeostasis, uncovering a previously unrecognised cellular axis underlying these host-adaptive traits. Together, these findings establish WD40-dependent PPI networks as key regulators of fungal pathogenicity and point to PPI-directed interventions as a potential antifungal strategy.

## Results

### Construction of the WD40 deletion and conditional knockdown mutants

The overall experimental workflow is illustrated in Fig. 1a. To identify WD40-encoding genes in the *C. neoformans* H99 genome, we used the WDSP database (http://www.wdspdb.com/wdspdb3/)^17^ and confirmed WD40 repeats through sequence analysis and domain classification with InterPro (https://www.ebi.ac.uk/interpro/). This approach identified 140 putative WD40 genes (Supplementary Data 1). Among these, 94 were classified as high-confidence candidates that contain at least six WD40 repeats, a feature required for stable β-propeller formation that underpins PPIs^18^ (Extended Data Fig. 1a, Fig. 1b). To assess the evolutionary conservation of these canonical WD40 proteins, we further performed comparative BLAST matrix analysis across representative fungal species, using a curated list of putative WD40 proteins identified in each species (Supplementary Data 2). This analysis showed that more than half are broadly conserved, whereas a subset is much less well conserved (Supplementary Data 3). We therefore focused subsequent analyses on these 94 canonical WD40 proteins.

**Fig. 1:**
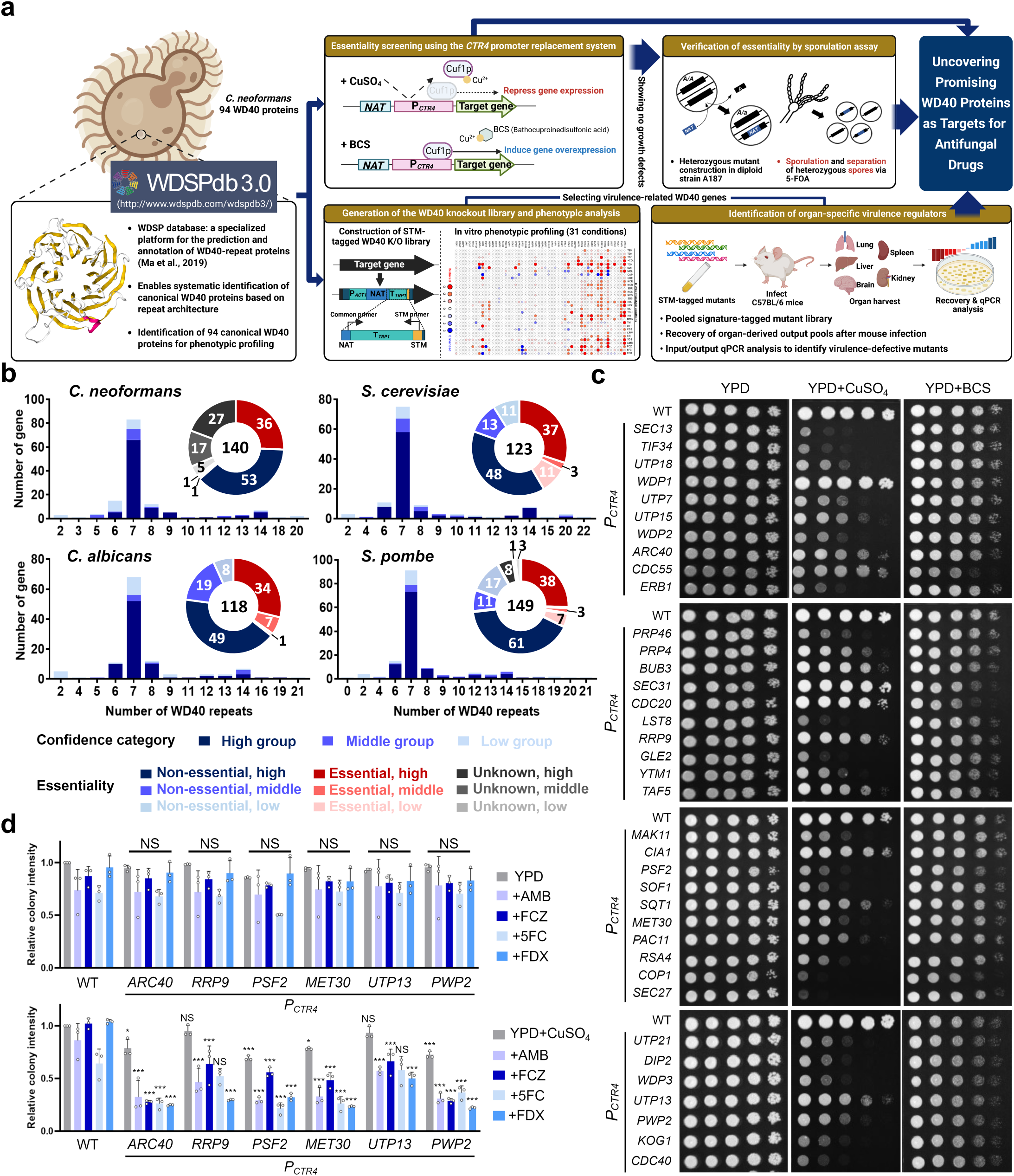
Conservation and essentiality analysis of 94 WD40 genes. **a,** Schematic overview of the identification and functional characterisation of WD40 proteins in *C. neoformans*. **b,** Variation in the number of WD40 repeats across fungal species, determined using the WDSP database (http://www.wdspdb.com/wdspdb3/), and essentiality of WD40 genes in key fungal species (Cn, Sc, Ca, and Sp). Essentiality in *C. neoformans* was assessed in this study, whereas data for *S. cerevisiae*, *C. albicans*, and *S. pombe* were obtained from the *Saccharomyces* Genome Database (http://www.yeastgenome.org), *Candida* Genome Database (http://www.candidagenome.org), and PomBase (http://www.pombase.org), respectively. **c**, Serial-dilution growth analysis of 37 *CTR4* promoter-replacement strains under basal conditions (YPD), suppression conditions (25 μM CuSO_4_), and overexpression conditions (200 μM BCS). Plates were incubated for 2 days and photographed. **d**, Quantification of relative colony intensity. Colony intensities were measured from 10^3^ dilution spots on day 4 and quantified using Image Lab 3.0 software. Mean values were normalised to the wild-type colony intensity under YPD (upper panel) or YPD + CuSO_4_ (lower panel) conditions. Relative colony intensity reflects the growth difference between WT and *CTR4* promoter replacement strains, providing a quantitative comparison of colony growth defects.

To investigate their pathobiological functions, we aimed to generate signature-tagged deletion mutants for all 94 canonical WD40 genes and to characterise their phenotypes in vitro and in vivo. Eleven WD40 genes had been partially characterised in previous studies, including *MSL1*^19^, *SWD1*^20^, *SWD2*, *SWD3*, *FAR8*^21^, *VPS15*^22^, *EED1*^23^, *GPB1*^24^, *GIB2*^25^, *TUP1*^26^ and *CDC4*^27^. However, signature-tagged mutants were available only for *VPS15*, *FAR8* and *MSL1*. For the remaining 91 WD40 genes, we sought to establish a high-quality mutant collection by generating at least two independent deletion strains per gene and verifying genotypes using diagnostic PCR and Southern blot analysis. This effort resulted in 98 newly constructed mutants representing 49 WD40 genes. In total, we analysed 103 mutant strains representing 52 WD40 genes. Detailed information on disruption strategies, primer sequences, Southern blot results, and phenotypic profiles is available in the *Cryptococcus neoformans* WD40 Phenome Database (http://WD40.cryptococcus.org/).

Despite repeated attempts, we were unable to generate knockout mutants for the remaining 42 WD40 genes, raising the possibility that these genes are essential for viability. To test this possibility, we replaced the native promoter of these genes with the copper-regulated *CTR4* promoter, which silences gene expression in the presence of copper^28^. This strategy successfully produced conditional knockdown strains for all but five genes (CNAG_06824, 06684, 01600, 07440 and 06772). Notably, 36 of the 37 promoter replacement strains exhibited clear growth defects under copper-repressive conditions (Fig. 1c), further supporting their essentiality. The sole exception was *WDP1*, whose conditional knockdown strain exhibited no detectable growth impairment under copper-repressive conditions, suggesting that *WDP1* may be dispensable for vegetative growth. To independently test this, we performed a diploid sporulation assay, in which recovery of viable haploid progeny carrying the disrupted allele indicates that the gene is non-essential for vegetative growth. Of 55 haploid spores analysed (34 *MAT***a** and 21 *MAT*α), 21 contained the knockout allele (15 *MAT***a** and 6 *MAT*α), confirming that *WDP1* is non-essential (Extended Data Fig. 2). Diagnostic PCR and independent reconstruction of the *wdp1*Δ in the H99 background further confirmed this conclusion.

Given the potential of essential WD40 proteins as antifungal drug targets, we next assessed whether their repression altered susceptibility to existing antifungal drugs. Repression of *UTP7, UTP15, ARC40, CDC55, ERB1, PRP46, BUB3, RRP9, TAF5, PSF2, MET30, PAC11, UTP21, DIP2, WDP3, UTP13, PWP2, KOG1* and *CDC40* increased susceptibility to certain antifungal drugs. Among these, repression of *ARC40, RRP9, PSF2, MET30, UTP13* and *PWP2* led to synergistic effects with multiple antifungal drugs, highlighting these genes as promising targets for combination therapy (Fig. 1d and Extended Data Fig. 1b). Notably, repression alone under basal conditions did not impair growth; however, when combined with sublethal antifungal concentrations, it led to a pronounced reduction in growth relative to controls. These findings underscore essential WD40 proteins, particularly those showing synergistic drug effects, as promising targets for antifungal therapy.

### Systematic in vitro and in vivo phenotypic profiling of *C. neoformans* WD40 proteins

Utilising the WD40 deletion mutant library, we conducted in vitro and in vivo phenotypic analyses as outlined in Fig. 2a, b. The complete phenome dataset monitored under 31 distinct in vitro conditions is presented as a heatmap (Fig. 2c; Supplementary Data 4). To evaluate their role in pathogenicity, we performed signature-tagged mutagenesis (STM)-based organ infectivity assays in a murine intranasal infection model (Fig. 2c, Supplementary Data 5). This analysis identified 31 WD40 genes required for infectivity across multiple organs. Several, including Swd1, Swd2, Swd3 (subunits of the COMPASS complex), Vps15, Tup1, Far8 and Msl1, have previously been established as virulence determinants, validating our approach.

**Fig. 2:**
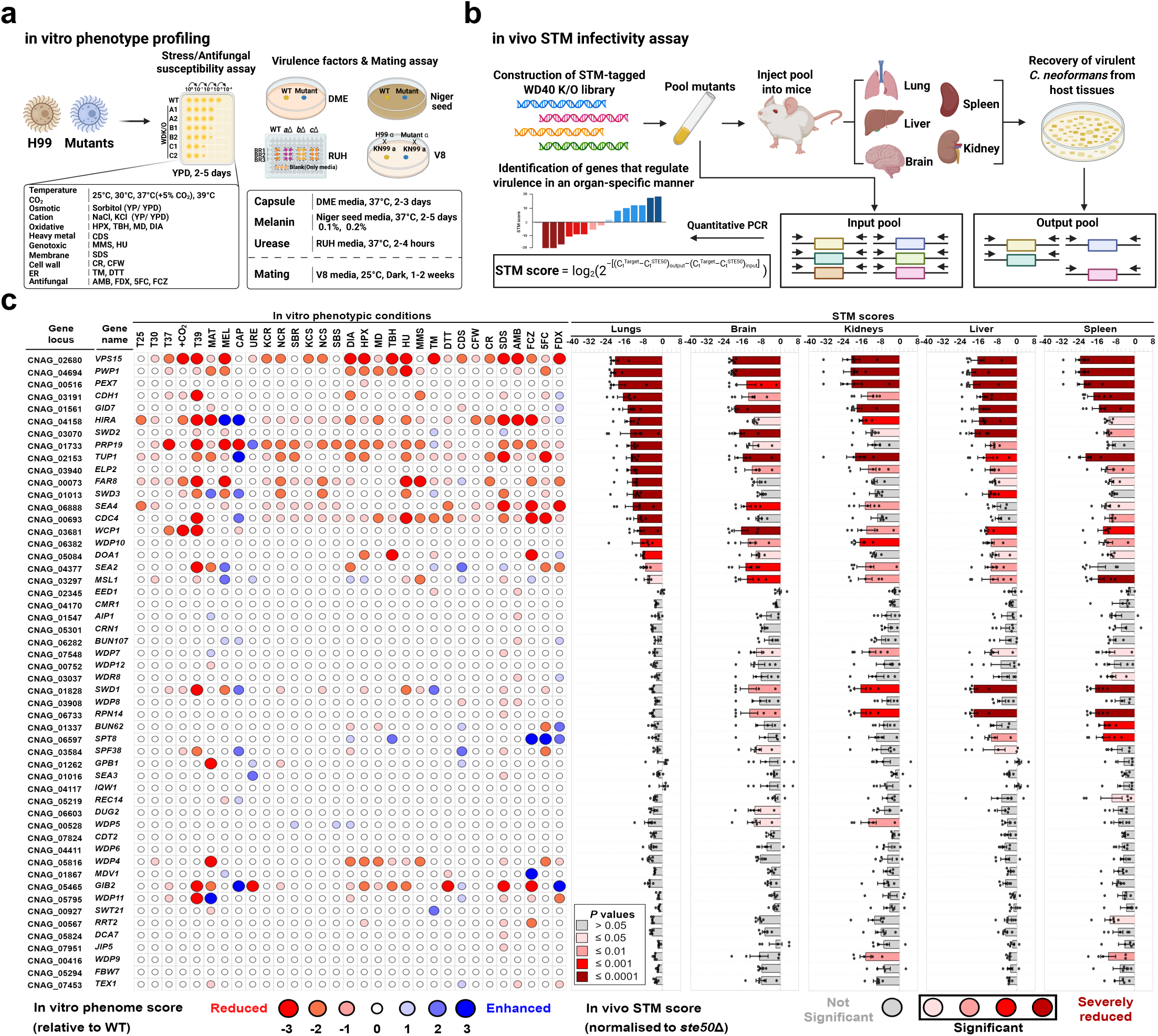
Systematic in vitro and in vivo phenotypic profiling of WD40 proteins in *C. neoformans*. **a**, Schematic overview of the in vitro phenotypic profiling strategy. **b**, Schematic overview of the STM-based murine infectivity assay. **c**, Integrated summary of in vitro phenotypic profiles across 31 conditions and organ-specific STM scores from the in vivo infectivity assay. In vitro phenotypic traits were assessed under 31 growth conditions and scored on a 7-point scale (-3, strongly reduced/susceptible; -2, moderately reduced/susceptible; -1, weakly reduced/susceptible; 0, wild-type like; +1, weakly enhanced/tolerant; +2, moderately enhanced/tolerant; +3, strongly enhanced/tolerant). In vivo infectivity was evaluated using an STM-based murine infectivity assay, with *ste50*Δ serving as a positive control for normalisation. Mice were infected intranasally with pooled WD40 mutants, and fungal burden was quantified in lungs, brain, kidney, liver and spleen. STM scores were calculated as log-transformed values, normalised to the *ste50*Δ mutant, and scored on a 5-point scale. All phenotypic and infectivity data are available in the *Cryptococcus neoformans* WD40 Phenome Database (https://WD40.cryptococcus.org/) More than three biologically independent experiments were performed for each phenotypic trait. **Abbreviations**: T25, 25°C; T30, 30°C; T37, 37°C; +CO_2_, 37°C + 5% CO_2_; T39, 39°C; CAP, capsule production; MEL, melanin production; URE, urease production; MAT, mating; HPX, hydrogen peroxide (3 to 3.5 mM); TBH, tert-butyl hydroperoxide (0.6 to 0.7 mM); MD, menadione (0.02 to 0.03 mM); DIA, diamide (2 to 2.5 mM); MMS, methyl methanesulfonate (0.03 to 0.04 %); HU, hydroxyurea (100 to 110 mM); 5FC, 5-flucytosine (300 to 500 μg/ml); AMB, amphotericin B (1.6 to 1.8 μg/ml); FCZ, fluconazole (10 to 13 μg/ml); FDX, fludioxonil (1 to 3 μg/ml); TM, tunicamycin (0.3 to 0.4 μg/ml); DTT, dithiothreitol (16 to 18 mM); CDS, cadmium sulphate (25 to 30 μM); SDS, sodium dodecyl sulphate (0.03 to 0.04 %); CR, Congo red (0.8 to 1 %); CFW, calcofluor white (3 to 5 mg/ml); KCR, YPD+1.5 M KCl; NCR, YPD+1.5 M NaCl; SBR, YPD+2 M sorbitol; KCS, YP+1 M KCl; NCS, YP+1 M NaCl; SBS, YP+2 M sorbitol. Heatmap scales for temperature-dependent growth, stress responses, and antifungal susceptibility were calculated relative to the T30 for each mutant. Abbreviations for stress conditions and colour codes for functional categorisation are detailed in the right panel.

Strikingly, mutants lacking *VPS15*, *PWP1*, *PEX7*, *CDH1*, *GID7*, *HIRA*, *SWD2*, *TUP1*, *ELP2*, *WCP1*, *WDP10*, *SEA2* or *MSL1* exhibited reduced infectivity across all five tested organs (lungs, brain, kidney, liver, and spleen). Many of these genes are central to production of major virulence factors (capsule, melanin and urease), stress adaptation and antifungal resistance. Notably, the *pwp1*Δ mutant displayed the most severe infectivity reduction across all tested organs, correlating with its increased sensitivity to high temperature, oxidative stress, and genotoxic stress and defective melanogenesis. By contrast, mutants of *PRP19*, *SWD3*, *FAR8*, *SEA4*, *CDC4*, *DOA1*, *WDP7*, *WDR8*, *SWD1*, *RPN14*, *BUN62*, *SPT8*, *SPF38*, *REC14*, *DUG2*, *WDP5*, *RRT2* and *WDP9* displayed reduced pathogenicity in at least one tested organ, though not consistently across all tissues. Thus, the STM analysis distinguished WD40 genes broadly required for in vivo fitness from those with organ-specific contributions, suggesting that distinct WD40-dependent pathways support cryptococcal adaptation to different host niches. In particular, several WD40-containing components of epigenetic complexes, including COMPASS, HIRA, SAGA, EED, ERP2 and TUP1, and post-transcriptional regulators, such as PRP19, played pleiotropic roles in diverse pathobiological traits of *C. neoformans.* Collectively, these results provide strong evidence that WD40 proteins are integral regulators of *C. neoformans* virulence.

### The WD40-containing cyclophilin promotes high CO_2_ tolerance in *C. neoformans*

In addition to classical cryptococcal virulence factors, CO_2_ tolerance has emerged as a key determinant of cryptococcal pathogenicity, reflecting the sharp difference in CO_2_ levels between the environment (∼0.04%) and the mammalian host (∼5%). Previous studies have implicated several signalling pathways in CO_2_ tolerance, including Ras, calcineurin, the Mpk1 MAPK signalling and the RAM (Regulator of Ace2 and Morphogenesis) pathway^29^. More recently, genetic loci linking CO_2_ tolerance to cryptococcal virulence have been identified^13^.

We found that several WD40 complexes also contributed to CO_2_ tolerance (Fig. 3a). Mutants lacking components of COMPASS (*swd1*Δ and *swd3*Δ), STRIPAK (*far8*Δ), HIRA (*hira*Δ), Vps34/PI3K (*vps15*Δ), U5 small nuclear ribonucleoprotein (*spf38*Δ), APC/C (*cdh1*Δ), and a cyclophilin (CNAG_03681) displayed impaired growth under 5% CO_2_ at 37 °C compared with ambient conditions. Among these, the CNAG_03681 mutant was most sensitive, comparable to RAM pathway mutants (Fig. 3a). Because CNAG_03681 encodes a protein with an N-terminal WD40 domain and a C-terminal cyclophilin-type peptidyl-prolyl cis-trans isomerase (PPIase) domain, we designated it Wcp1 (<u>W</u>D40-containing <u>c</u>yclo<u>p</u>hilin 1) (Fig. 3b).

**Fig. 3:**
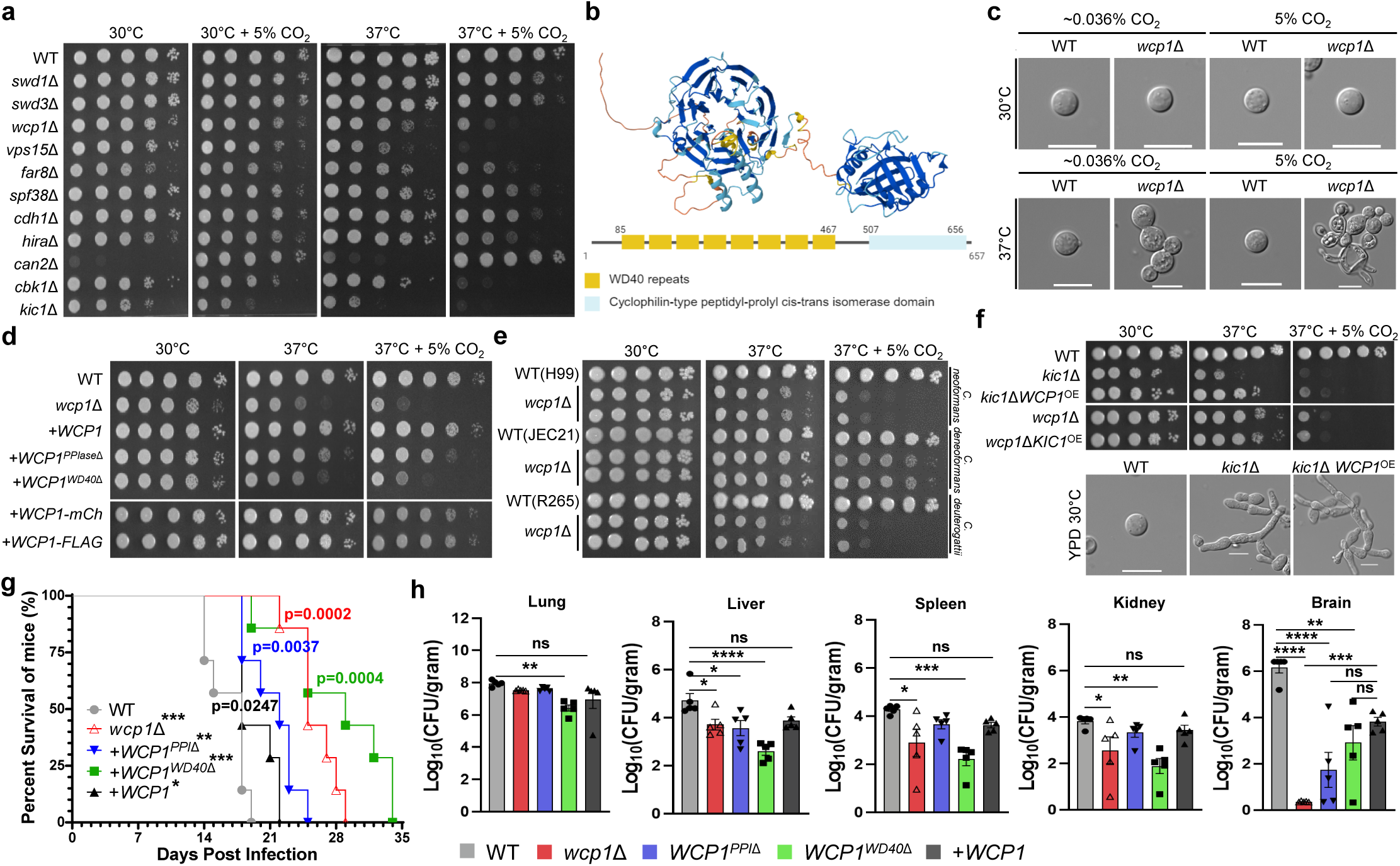
WD40-containing cyclophilin critical for high CO_2_ tolerance. **a,** Screening of WD40 genes involved in CO_2_ tolerance using a library of 52 knockout mutants. Serial-dilution growth analysis was performed under 30 °C with 5% CO_2_ and 37 °C with 5% CO_2_. Plates were photographed on day 2 (30 °C) and day 3 (37 °C). **b,** Predicted protein and domain structure of Wcp1. Protein structure was modelled using AlphaFold, and domain information was retrieved from InterPro (https://www.ebi.ac.uk/interpro/), UniProt (https://www.uniprot.org/), and WDSPdb 3.0 (http://www.wdspdb.com/wdspdb3/). **c,** Morphological analysis of *wcp1*Δ under high-temperature and CO_2_ conditions. Wild-type and *wcp1*Δ strains were cultured overnight, spotted on YPD, and incubated at 30 °C or 37 °C in ambient or 5% CO_2_ conditions for 2 days. Colonies were resuspended in PBS and observed microscopically (scale bar, 10 μm). **d,** Functional analysis of Wcp1 by complementation with non-tagged, mCherry-tagged and FLAG-tagged alleles, or with domain-deletion alleles lacking either the PPIase or WD40 domain. Plates were photographed on day 2. **e,** Conserved role of Wcp1 across *Cryptococcus* strains. *wcp1*Δ mutants in *C. neoformans* (H99), *C. deneoformans* (JEC21), and *C. deuterogattii* (R265) were assayed for CO_2_ tolerance at 37 °C and 37 °C with 5% CO_2_. Plates were photographed on day 3. **f,** Epistasis analysis between Wcp1 and Kic1 (RAM pathway). Overexpression of *WCP1* in the *kic1*Δ background and overexpression of *KIC1* in the *wcp1*Δ background were assessed for growth phenotypes. Plates were photographed on day 3. **g**, Survival assay in the BALB/c murine infection model (n = 7 per group). Strains tested were WT (H99), *wcp1*Δ, +*WCP1*, +*WCP1^PPIase^*^Δ^ and +*WCP1^WD40^*^Δ^. Statistical differences were assessed using the log-rank (Mantel–Cox) test. **h**, Fungal burden assay at 18 dpi (n = 5 per group). A separate cohort of mice was euthanised at 18 days post-infection (dpi), and organ fungal loads were quantified as CFU per gram of tissue. For each organ, statistical significance was assessed by one-way ANOVA followed by Tukey’s multiple-comparisons test.

The *wcp1*Δ mutant showed abnormal morphogenesis under 5% CO_2_ at 37 °C, including cell aggregation and vacuolar enlargement (Fig. 3c). These defects were restored in both the untagged complemented strain (*wcp1*Δ::*WCP1*) and the tagged complemented strains expressing *WCP1*-*mCherry* and *WCP1*-*4×FLAG* (Fig. 3d, Extended Data Fig. 3), confirming that the tagged fusion proteins are functional. *C. neoformans* encodes 13 cyclophilins, but Wcp1 is the only member harbouring a WD40 domain, containing eight WD40 repeats and one PPIase domain (Fig. 3b). Domain-deletion complementation revealed that both domains are required for full function, with loss of the WD40 domain causing more severe CO_2_- and temperature-sensitive growth defects than loss of the PPIase domain. Thus, Wcp1 function relies on both PPIase enzymatic activity and WD40-mediated interactions. Importantly, orthologues in *Cryptococcus deneoformans* and *Cryptococcus deuterogattii* were also essential for CO_2_/heat tolerance (Fig. 3e), indicating that this function is conserved across the pathogenic *Cryptococcus* species complex.

To test genetic interactions between *WCP1* and the RAM pathway, we used *kic1*Δ as a representative RAM-pathway mutant, as Kic1 is a core RAM kinase required for cryptococcal morphogenesis, thermotolerance and CO_2_ adaptation. Overexpression of *WCP1* partially rescued the thermosensitivity of the *kic1*Δ mutant but not its CO_2_ sensitivity, whereas *KIC1* overexpression failed to complement *wcp1*Δ (Fig. 3f). These results suggest that Wcp1 may function downstream of, or in parallel to, the RAM pathway, and that thermotolerance and CO_2_ tolerance are mediated by distinct mechanisms. The persistence of pseudohyphal morphology in the *kic1*Δ *WCP1^OE^*strain further supports separate regulatory circuits for morphology, temperature adaptation, and CO_2_ responses.

We next assessed the contribution of Wcp1 to *C. neoformans* virulence in a murine model of systemic cryptococcosis. Mice infected with the *wcp1*Δ mutant or the WD40-domain-deletion strain showed markedly prolonged survival and substantially reduced mortality, indicating that loss of Wcp1 or its WD40 scaffold profoundly attenuates virulence (Fig. 3g). By contrast, the PPIase-domain–deletion strain exhibited a modest reduction in virulence relative to the full-length *WCP1*-complemented strain, which caused rapid mortality consistent with full virulence. Fungal burden analyses at 18 days post-infection, corresponding approximately to the median survival time of mice infected with the wild-type strain, further revealed organ-specific defects in dissemination and CNS invasion (Fig. 3h). Lung CFU were comparable across strains, indicating that Wcp1 is largely dispensable for initial pulmonary colonisation. However, fungal burdens in the liver, spleen and kidney were substantially reduced in mice infected with the *wcp1*Δ and WD40-domain–deletion strains, whereas the PPIase-domain–deletion strain and the full-length *WCP1*-complemented strain reached near wild-type levels in these peripheral organs. The most pronounced differences were observed in the brain: fungal loads were substantially reduced in the *wcp1*Δ mutant, and were lower in the PPIase-domain–deletion strain than in the WD40-domain–deletion strain, suggesting that the PPIase domain makes a particularly important contribution to CNS invasion, while the WD40 scaffold plays a secondary yet significant role. Together, these data indicate that Wcp1 is critical for systemic dissemination and CNS infection in the mammalian host, with its PPIase and WD40 domains contributing distinctly to full virulence.

### Wcp1 contributes to CO_2_/heat tolerance via acidic pH adaptation

To determine whether Wcp1 contributes to virulence only under mammalian host-like conditions, we used a *Drosophila melanogaster* infection model maintained at 30 °C in ambient atmospheric CO_2_ (∼0.04%) (Fig. 4a). Because this system does not recapitulate the 37 °C/5% CO_2_ environment of mammalian hosts, it enabled us to test whether the virulence defect of the *wcp1*Δ mutant persists outside host-like thermal and CO_2_ stress conditions. In this model, flies infected with the *wcp1*Δ mutant or with a strain lacking the PPIase domain showed markedly increased survival compared with those infected with the wild-type strain (Fig. 4b). The persistence of this attenuation under moderate-temperature, low-CO_2_ conditions indicates that Wcp1 promotes virulence through mechanisms that cannot be explained solely by thermotolerance or CO_2_ tolerance, suggesting an additional role in fundamental aspects of host adaptation.

**Fig. 4:**
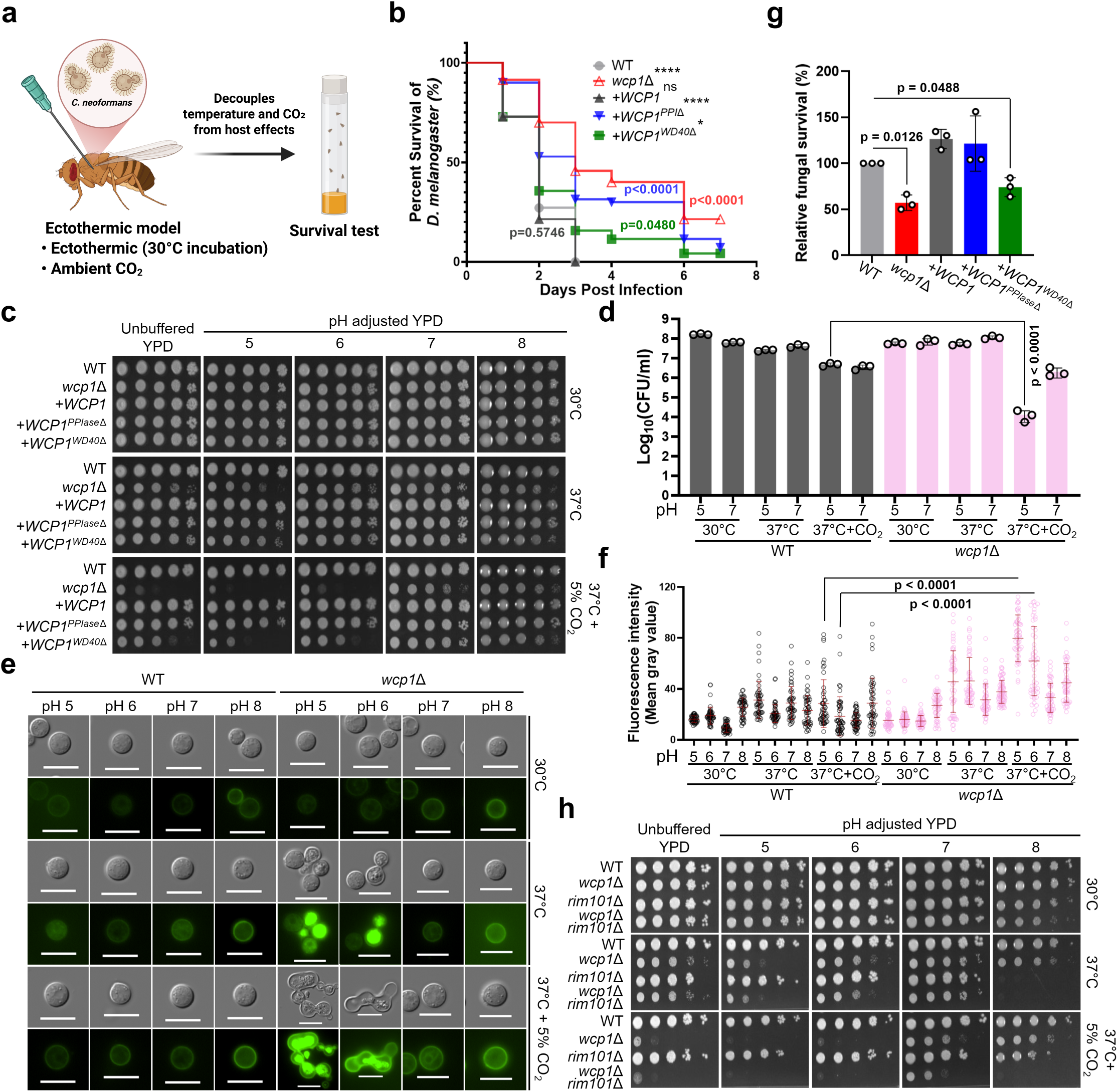
Wcp1 links CO_2_/heat tolerance to acidic pH adaptation and intracellular acidification. **a**, Schematic of the *Drosophila melanogaster* infection model used to assess Wcp1-dependent virulence independently of host-like temperature and CO_2_ conditions. **b**, Survival of flies infected with WT, *wcp1*Δ, +*WCP1*, +*WCP1^PPIase^*^Δ^ and +*WCP1^WD40^*^Δ^ strains. For each strain, 50–70 flies were infected and divided into vials of 8–10 flies. Survival was monitored daily for 6 days at 30 °C and analysed using Kaplan–Meier curves with the log-rank test. **c**, pH-dependent growth analysis of WT, *wcp1*Δ, +*WCP1*, +*WCP1^PPIase^*^Δ^ and +*WCP1^WD40^*^Δ^ strains on unbuffered or pH-adjusted YPD under 30 °C, 37 °C and 37 °C + 5% CO_2_ conditions. Plates were photographed after 4 days of incubation. **d**, Colony-forming units (CFU) of WT and *wcp1*Δ under the indicated pH, temperature and CO_2_ conditions. Overnight cultures were subcultured into fresh YPD, grown to OD_600_ 0.8, divided into pH 5 and pH 7 conditions, and incubated for 2 days at 30 °C, 37 °C or 37 °C + 5% CO_2_. Cells were then harvested, serially diluted and plated on YPD agar containing chloramphenicol (100 μg/ml) for CFU determination. Data are from three biologically independent experiments. **e**, Representative brightfield and LysoSensor fluorescence images of WT and *wcp1*Δ cells under the indicated pH, temperature and CO_2_ conditions. **f**, Quantification of LysoSensor fluorescence intensity shown in e. At least 50 cells were quantified per strain and condition. Data are mean ± SD (standard deviation); statistical differences within each condition were calculated using ordinary one-way ANOVA followed by Tukey’s multiple-comparisons test. **g**, Relative fungal survival within macrophages for the indicated strains. Data are mean ± SD from three biological replicates; statistical differences were calculated using a one-sample t-test against the normalised WT value of 100%. **h**, Genetic interaction analysis between WT, *wcp1*Δ, *rim101*Δ and *wcp1*Δ *rim101*Δ under unbuffered or pH-adjusted YPD conditions.

CO_2_ freely diffuses into cells and rapidly equilibrates with bicarbonate (HCO) and protons (H) through carbonic anhydrase-catalysed and spontaneous hydration reactions^30,31^. Consequently, elevated CO_2_ can transiently acidify the cytosol. High temperature can further exacerbate intracellular acidification by boosting metabolic and respiratory flux, compromising membrane integrity and proton gradients, and depleting ATP required for proton extrusion and buffering, thereby lowering cytosolic pH^31–33^. We therefore compared in vitro growth of wild-type and *wcp1*Δ strains under elevated temperature and CO_2_ across a series of buffered pH conditions using YPD medium (Fig. 4c), with parallel experiments performed in RPMI medium (Extended Data Fig. 4a). Under combined high temperature and CO_2_, the *wcp1*Δ mutant showed severe growth inhibition at acidic pH, whereas growth was substantially but incompletely rescued at alkaline conditions. Domain-deletion strains revealed that deletion of the WD40 domain largely recapitulated the acidic pH defect, indicating a pivotal role for the WD40 scaffold in adaptation to acidic environments under high temperature and CO_2_. Because this defect was slightly less severe than that of the *wcp1*Δ mutant, the PPIase domain likely provides an additional, modest contribution. Consistent with the serial-dilution growth analysis, CFU quantification showed that the *wcp1*Δ strain had markedly reduced survival under acidic conditions, whereas this defect was largely alleviated at higher pH (Fig. 4d, Extended Data Fig. 4b).

To further validate these phenotypes, we quantified intracellular acidification using the acidotropic, pH-sensitive fluorescent probe LysoSensor (Fig. 4e, f, Extended Data Fig. 4c). At 37 °C and pH 5, the *wcp1*Δ mutant displayed markedly higher LysoSensor fluorescence than the wild type, indicating enhanced intracellular acidification. This difference increased further at 37 °C with 5% CO_2_, particularly at pH 5 and pH 6, where high CO_2_ consistently induced stronger acidification than ambient CO_2_. We next assessed whether this defect compromises survival within macrophages, where the phagolysosome imposes a highly acidic microenvironment. Consistent with this hypothesis, macrophage killing assays showed significantly reduced survival of the *wcp1*Δ and WD40-domain–deletion strains relative to the wild type (Fig. 4g). Together, these data identify Wcp1 as a key determinant of adaptation to acidic environments under elevated temperature and CO_2_, supporting survival in host-like niches such as macrophages.

We next examined the relationship between Wcp1 and the Rim101-mediated pH adaptation pathways. Rim101 is a conserved fungal transcription factor required for alkaline pH adaptation^34^, providing a contrasting genetic reference to the acidic pH growth defect of *wcp1*Δ. Consistent with this role, the *rim101*Δ mutant showed severe growth defects at 37 °C under alkaline, but not acidic, conditions, underscoring the essential role of Rim101 in alkaline adaptation. Notably, the growth defect of *rim101*Δ at 37 °C was substantially alleviated by 5% CO_2_ (Fig. 4h), consistent with CO_2_-driven intracellular acidification partially counteracting alkaline stress. Although the *wcp1*Δ mutant grew better at alkaline than acidic pH, the *wcp1*Δ *rim101*Δ double mutant failed to grow at all under alkaline conditions, indicating that Wcp1 and Rim101 provide non-redundant functions required for survival in alkaline environments (Fig. 4h). Overexpressing *WCP1* in *rim101*Δ or *RIM101* in *wcp1*Δ did not rescue the respective mutant phenotypes (Extended Data Fig. 5), suggesting that increased expression of one pathway cannot compensate for loss of the other. Together, these results support distinct yet complementary roles for Wcp1 and Rim101: Wcp1 is critical for adaptation to acidic pH conditions, whereas both are required for growth under alkaline conditions at host physiological temperature. Thus, Wcp1 and Rim101 define complementary arms of cryptococcal pH homeostasis, with Wcp1 buffering acidification stress and Rim101 driving alkaline adaptation.

### Wcp1 forms a pH-dependent interactome that supports adaptation to acidification stress

To define the Wcp1-associated protein network, we performed affinity purification–mass spectrometry (AP–MS) using functionally validated Wcp1-4×FLAG and Wcp1-mCherry strains at pH 5 and pH 7 at 37 °C (Fig. 5a; Extended Data Fig. 3a,b). Across all conditions, we identified a total of 191 Wcp1-associated proteins. Comparative analysis revealed a markedly pH-dependent interaction landscape (Fig. 5b, Supplementary Data 6). Specifically, 108 proteins were detected exclusively under acidic conditions (pH 5), whereas 77 proteins were shared between conditions, and only 6 proteins were uniquely detected at pH 7, indicating that Wcp1 preferentially assembles protein complexes under acid stress.

**Fig. 5:**
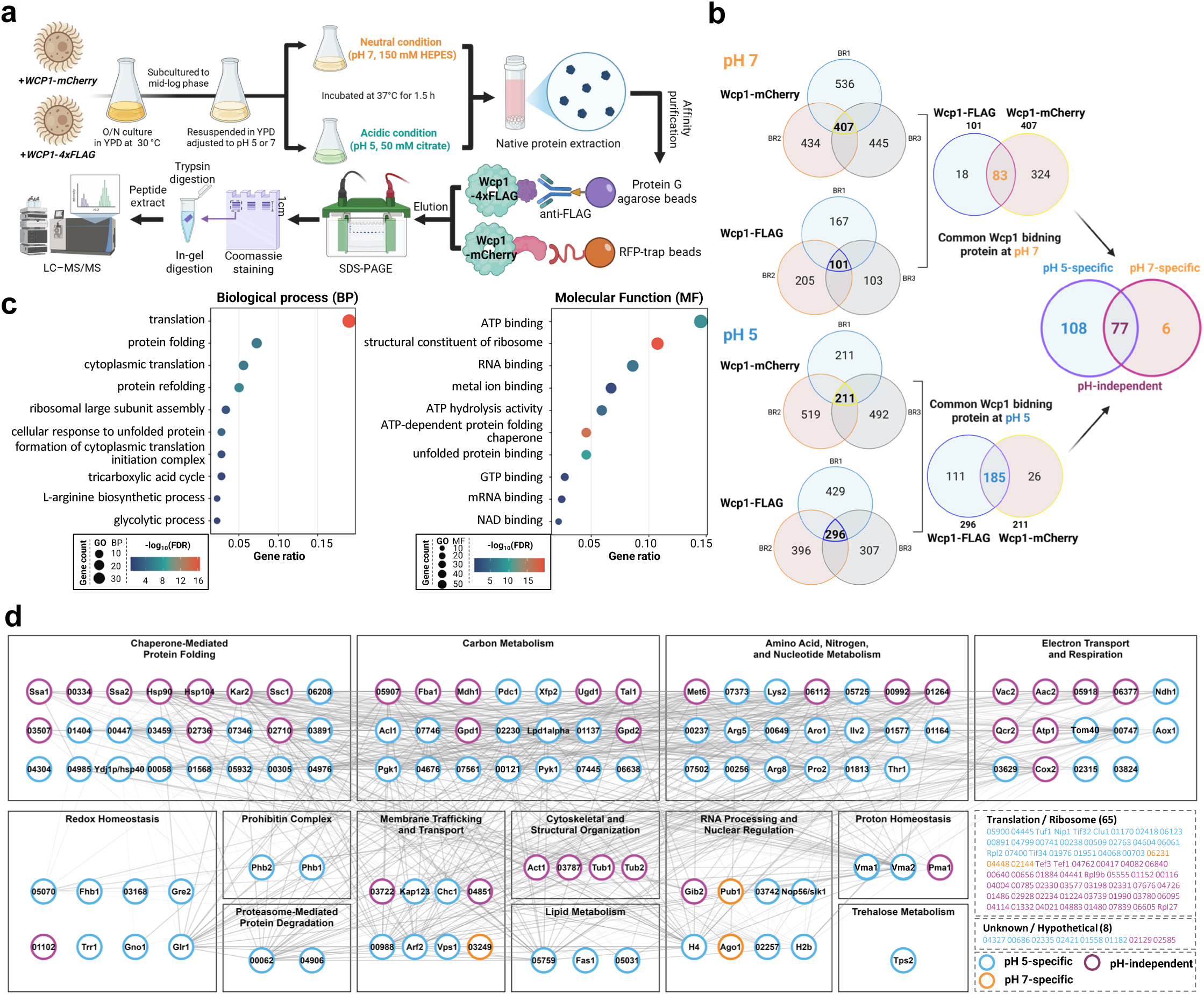
Wcp1 forms a pH-dependent interactome under acidification stress. **a**, Experimental workflow for affinity purification–mass spectrometry analysis of Wcp1-associated proteins at pH 5 and pH 7. **b**, Venn diagram showing pH 5-specific, pH 7-specific and shared Wcp1-associated proteins identified by AP–MS using Wcp1-mCherry and Wcp1-4×FLAG strains. **c**, Gene Ontology enrichment analysis of the Wcp1 interactome, showing over-represented biological process and molecular function categories. **d**, STRING-based protein interaction network of the Wcp1 interactome. This was visualised using Cytoscape v3.10.3.

Gene ontology (GO) enrichment analysis of the Wcp1 interactome showed over-representation of biological processes related to translation, protein folding and central metabolism, including the tricarboxylic acid cycle, glycolysis and amino acid biosynthesis. Enriched molecular function terms included ATP binding, ATP hydrolysis activity, chaperone activity and RNA binding (Fig. 5c), linking the Wcp1 interactome to proteostasis, translational machinery and core metabolic pathways. To examine the organisation of these interactions, we constructed a STRING-based interaction network. This analysis resolved the Wcp1 interactome into distinct functional modules, including chaperone-mediated protein folding, carbon metabolism, amino acid and nucleotide metabolism, electron transport and respiration, and proton and redox homeostasis (Fig. 5d). Notably, the interactome included direct pH homeostasis factors, including V-ATPase subunits (CNAG_02326 and CNAG_04439), which mediate vacuolar proton transport, and Pma1, the plasma membrane H^+^-ATPase, indicating that Wcp1 is physically coupled to core proton-handling machinery under acidic conditions.

Beyond this direct pH homeostasis axis, the Wcp1 interactome pointed to two additional cellular processes supporting acidification defence. First, the enrichment of chaperones and redox-regulating proteins suggested that Wcp1 may mitigate proteostatic and oxidative stress that accompany acidification under elevated CO_2_ and temperature. Consistent with this prediction, the *wcp1*Δ mutant was hypersensitive to ER and oxidative stress at elevated temperature (Extended Data Fig. 6a). Second, the enrichment of metabolic and respiratory proteins suggested that Wcp1 may sustain the energetic supply required to fuel ATP-dependent pH homeostasis machinery. To test this, we examined growth on glucose and acetate. Wild-type cells grew comparably on both carbon sources even under combined acidic, CO_2_ and heat stress, whereas the *wcp1*Δ mutant displayed a marked growth defect on acetate under these conditions (Extended Data Fig. 6b). Because acetate catabolism requires mitochondrial respiration, this phenotype supports a role for Wcp1 in sustaining respiratory metabolism under acidification stress. Together, these data indicate that Wcp1 coordinates a multi-layered defence against acidification stress, integrating proton-handling, proteostasis, oxidative stress defence and respiratory metabolism.

### Wcp1 regulates a pH-dependent transcriptional programme for adaptation to acidification stress

Having defined a pH-dependent Wcp1 interactome integrating proton handling, proteostasis and metabolism, we next asked whether Wcp1 also contributes to transcriptional responses. Wcp1–mCherry showed predominant nuclear localisation across all tested conditions (30°C, 37°C and 37°C with 5% CO_2_), and AP–MS identified interactions with chromatin components, including the core histones H4 and H2B (Fig. 6a). To address this, we performed RNA-seq analysis under neutral and acidic conditions. Principal component analysis showed clear separation by genotype and condition, with tight clustering of biological replicates, and revealed a pronounced Wcp1-dependent shift that was most evident under acidic conditions (Extended Data Fig. 7a). Differential expression analysis revealed extensive transcriptome remodelling in the absence of Wcp1. At neutral pH, deletion of *WCP1* altered the expression of 1,104 genes relative to the wild type (759 upregulated and 345 downregulated), whereas under acidic conditions this increased to 1,293 genes (1,139 upregulated and 154 downregulated). These data indicate not only a greater regulatory impact of Wcp1 under acid stress, but also a marked shift towards gene upregulation in its absence. Consistent with this, the transcriptional response to acidification was balanced in the wild type but strongly skewed towards gene upregulation in the *wcp1*Δ mutant, indicating that Wcp1 constrains and organises the acid-responsive transcriptional programme (Fig. 6b, Supplementary Data 7). This bias was further reflected in gene composition: whereas the downregulated gene sets were relatively enriched for functionally annotated genes, the strongly expanded upregulated gene sets in *wcp1*Δ contained a large proportion of functionally unannotated transcripts, including hypothetical and uncharacterised proteins, particularly under acidic conditions and during acidification responses (Extended Data Fig. 7c). These patterns suggest that loss of Wcp1 not only amplifies transcriptional output under acid stress but also derepresses a poorly coordinated gene-expression programme enriched in functionally unresolved loci.

**Fig. 6:**
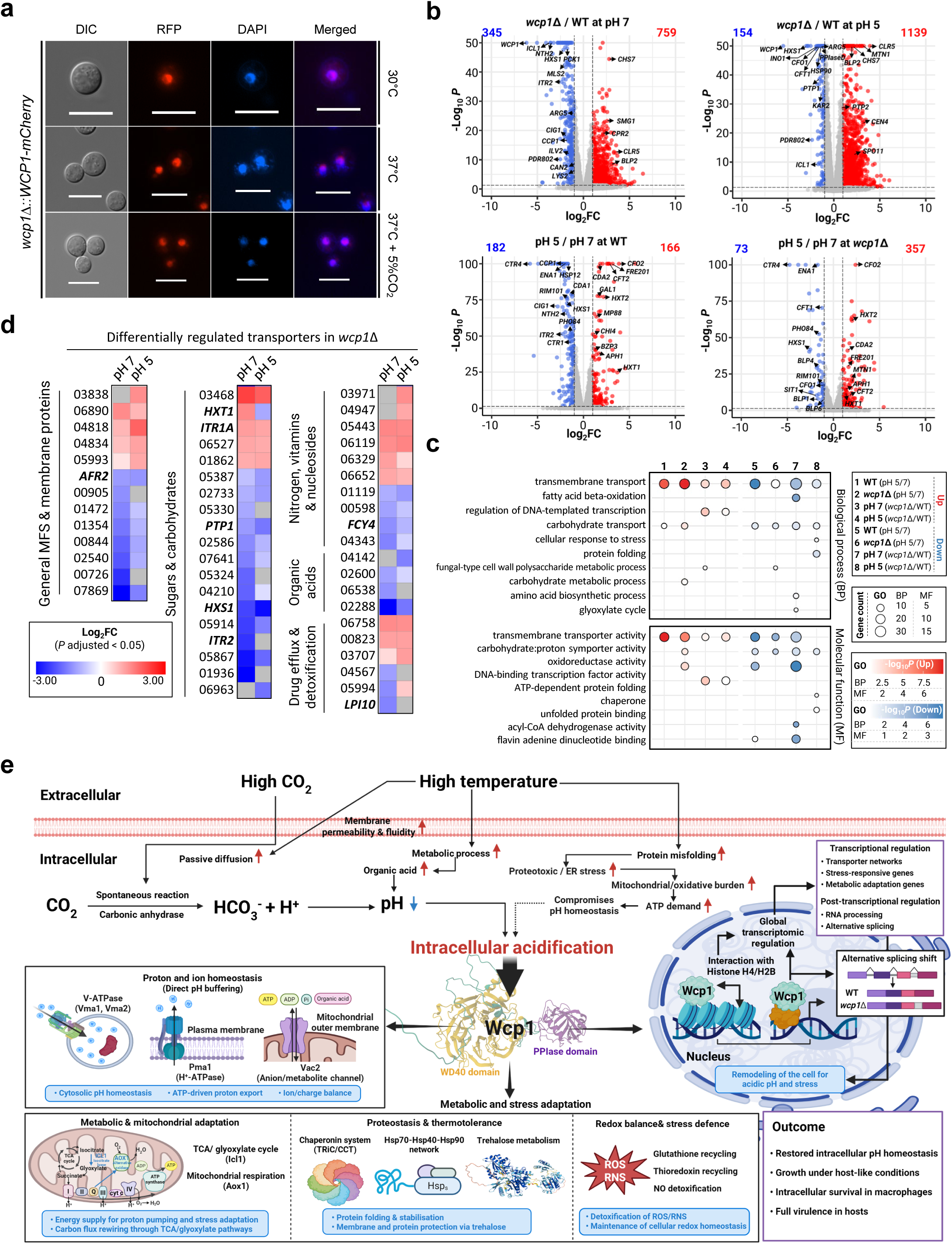
Wcp1 regulates a pH-dependent transcriptional programme for adaptation to acidification stress. **a,** Fluorescence microscopy of the Wcp1-mCherry strain under 30 °C, 37 °C and 37 °C + 5% CO_2_. **b**, Summary of Wcp1-dependent transcriptional changes at pH 7 and pH 5, including the relative proportions of upregulated and downregulated genes and the gene composition of each set. **c**, GO enrichment analysis of differentially expressed genes across the indicated transcriptomic comparisons. **d**, Differentially regulated transporter genes in *wcp1*Δ at pH 5 and pH 7, grouped into functional classes including general MFS and membrane proteins, sugars and carbohydrates, nitrogen, vitamins and nucleosides, organic acids, and drug efflux and detoxification. **e**, Model of Wcp1-mediated adaptation to intracellular acidification under host-like CO_2_ and temperature stress.

Because nuclear localisation and chromatin association raised the possibility that Wcp1 influences RNA regulation more broadly, we next examined alternative splicing. Acidification alone altered relatively few splicing events in either the wild type or the *wcp1*Δ mutant. By contrast, comparison between the two genotypes revealed substantially more alternative splicing differences at both neutral and acidic pH (Extended Data Fig. 7d), indicating that Wcp1 affects RNA processing beyond its effects on steady-state transcript abundance. A representative example was observed in CNAG_01295, which encodes an acyl carrier protein and showed a pronounced shift in splice-junction usage in the *wcp1*Δ mutant under acidic conditions (Extended Data Fig. 7e). Together, these findings suggest that Wcp1 organises the acidification-response programme at both transcriptional and post-transcriptional levels.

Consistent with this dual role, GO enrichment analysis of Wcp1-dependent genes revealed processes directly linked to pH homeostasis and stress buffering, including transmembrane transport, regulation of DNA-templated transcription, protein folding and cellular stress responses, alongside broad metabolic rewiring affecting carbohydrate and amino-acid metabolism, lipid catabolism and cell-wall polysaccharide pathways (Fig. 6c, Extended Data Fig. 7b). Further supporting a central role in ionic and pH homeostasis, transporter-focused analyses revealed coordinated, pH-dependent dysregulation of multiple membrane transporters in *wcp1*Δ, including major facilitator superfamily (MFS) transporters, nutrient transporters (*HXT1*, *ITR1a/ITR2*, *HXS1*), nitrogen, vitamin, nucleoside transporters such as *FCY4*, organic-acid transporters, and drug efflux/detoxification modules (Fig. 6d). Together with the proteomic evidence that Wcp1 associates with core proton-handling machinery, these transcriptional changes indicate that Wcp1 promotes adaptation to acidic pH not only through direct interaction with pH homeostasis factors, but also through coordinated regulation of membrane transporter expression.

## Discussion

PPIs underpin most biological processes, yet genome-wide functional studies of PPI scaffolds in fungal pathogens remain limited. Here we systematically surveyed 94 canonical WD40 proteins in *C. neoformans*, generating 98 mutants and phenotypically profiling 103 strains representing 52 WD40 proteins. Conditional knockdown indicated that 36 of the remaining WD40 genes are likely essential, consistent with recent TnSeq data confirming essentiality for 35 of these (except Bub3)^35^. Together with five genes refractory to deletion or knockdown—three of which (*PAC1*, *IPI3* and *RRB1*) have been independently validated as essential^35^—we estimate that 39 of 94 WD40 proteins (42%) are essential, comparable to *S. cerevisiae* (44%), *C. albicans* (42%) and *S. pombe* (36%). Cross-species comparison defines a core set of 22 essential fungal WD40 proteins (Sqt1, Sof1, Erb1, Sec13, Sec27, Utp7, Utp13, Utp15, Utp18, Utp21, Dip2, Arc40, Pwp2, Prp4, Prp46, Tif34, Taf5, Psf2, Rrp9, Ytm1, Rsa4 and Kog1), with Bub3 uniquely essential in *C. neoformans*. Notably, repression of Arc40 (Arpc1), a component of the ARP2/3 actin polymerisation complex^36^, markedly increased susceptibility to amphotericin B, fluconazole and flucytosine, identifying the ARP2/3 complex as a promising target for combination antifungal therapy.

Beyond essentiality, our survey reveals that WD40 proteins shape a broad spectrum of pathobiological traits. The three major virulence factors—melanin, capsule and thermotolerance—were influenced by multiple WD40-containing complexes, including the previously characterised COMPASS, STRIPAK, TUP1 and PI3K-III complexes^20,21,26,37^, and previously unrecognised contributions from HIRA, PRP19, SEACAT and PWP1. These findings establish WD40 proteins as active organisers of chromatin-associated, post-transcriptional, vesicular and metabolic regulatory networks underlying cryptococcal pathogenicity.

A particularly striking pathobiological axis defined by our screen is CO_2_ tolerance. The mammalian host environment (∼5% CO_2_) differs sharply from ambient air (∼0.04%), but our data suggest that its impact is best understood as part of a coupled host-like stress state rather than as an isolated CO_2_ signal. Elevated CO_2_ and high temperature likely impose a combined intracellular acidification burden: CO_2_ hydration generates additional proton load, while heat increases CO_2_ diffusion rates and exacerbates membrane stress, proteotoxicity and the energetic cost of maintaining ion and proton gradients. CO_2_ tolerance can therefore be viewed as the ability to withstand a composite acidification-centred stress state (Fig. 6e). Whereas carbonic anhydrases are essential in ambient air but largely dispensable under host-like CO ^38^, we identify multiple WD40-containing complexes—COMPASS (Swd1/Swd3), PI3K-III (Vps15), U5 snRNP (Spf38), APC/C (Cdh1), HIRA (Hira), STRIPAK (Far8) and Wcp1—as critical determinants of CO_2_ adaptation. The functional breadth of these complexes underscores the multifactorial nature of CO_2_ tolerance, and their consistent dependence on 37 °C highlights a tight interplay between host temperature and CO_2_ stress.

Among these, Wcp1 stands out as a previously uncharacterised WD40-containing cyclophilin in which both the WD40 scaffold and the cyclophilin-type PPIase domain are required for function. Its role in CO_2_ adaptation is conserved across pathogenic *Cryptococcus* species. *wcp1*Δ mutants in the clinical *C. neoformans* H99 and *C. deuterogattii* R265 backgrounds displayed pronounced CO_2_ sensitivity at 37 °C, whereas the less virulent *C. deneoformans* JEC21 background showed only a mild defect, suggesting lineage-specific refinement of Wcp1 function in highly virulent species. Because PPIases are pharmacologically tractable and contribute to microbial pathogenicity^39,40^, the dual WD40–cyclophilin architecture of Wcp1 may represent a distinctive host-adaptation module within the pathogenic *Cryptococcus* species complex^41^.

Integrated proteomic and transcriptomic analyses support a working model in which Wcp1 acts through two coupled tiers of regulation (Fig. 6e). At the protein level, Wcp1 organises a pH-biased interactome connecting proton-handling machinery (V-ATPase and Pma1) with mitochondrial respiration, proteostasis, redox control and vesicular trafficking, enabling rapid reorganisation of protein complexes to mitigate acute acidification stress. In parallel, Wcp1 shapes a broader acid-responsive transcriptional programme—enriched for membrane transporters, stress-response and metabolic genes—that sustains longer-term pH homeostasis. The WD40 scaffold likely organises stress-responsive protein assemblies, whereas the cyclophilin-type PPIase domain may modulate conformational or post-translational states of client proteins.

This dual mode coherently explains otherwise disconnected phenotypes: acidic-pH-specific recruitment of TRiC/CCT and Hsp40/60/70 chaperones, with Wcp1-dependent folding gene expression points to proteostasis reinforcement under combined heat-acid stress; pH 5-specific interaction with Tps2 alongside Wcp1-dependent repression of *NTH2* suggests trehalose accumulation through coupled synthesis-degradation control; and the acidic-pH-specific association with Aox1, respiratory-chain components and prohibitins, together with the acetate growth defect of *wcp1*Δ, indicates that Wcp1 sustains the mitochondrial capacity needed to meet the ATP demand of proton extrusion and vacuolar sequestration. Wcp1-dependent expression of *CAN2*, *ICL1*, acid-responsive iron uptake genes and *PDR802* further extends Wcp1 function into CO_2_ sensing, alternative carbon utilisation, nutrient acquisition and virulence-associated remodelling. Together, these observations raise the possibility that acidic pH functions not merely as one stress among many, but as an integrative signal through which Wcp1 primes *C. neoformans* for the thermal, respiratory, oxidative and nutritional challenges of the host environment. This multi-layered buffering model may also explain the pronounced dissemination and CNS colonisation defects of *wcp1*Δ in vivo, where fungal cells must sustain pH homeostasis while encountering changing thermal, oxidative, respiratory and nutritional stresses across tissue niches. Within this framework, the genetic interaction with the RAM pathway and Rim101 are best interpreted as complementary regulatory inputs rather than the primary organising axis: *WCP1* overexpression partially rescued RAM-associated thermosensitivity, while interaction with Rim101 points to complementary arms of cryptococcal pH homeostasis. Future phosphoproteomic and graded pH/CO_2_ transcriptomic analyses will help resolve how these pathways intersect mechanistically.

In summary, our study identifies Wcp1 as a dual-capacity WD40–cyclophilin regulator that integrates pH-dependent protein-complex assembly with transcriptional adaptation. This architecture explains how *C. neoformans* coordinates proton handling, mitochondrial metabolism, proteostasis, transporter regulation and virulence-associated functions under host-relevant CO_2_ and temperature stress. More broadly, our work establishes WD40-dependent PPI networks as central regulators of fungal pathogenicity and highlights PPI scaffolds as an underexplored class of antifungal targets.

## Methods

### Ethics statement

Animal care and all experiments were conducted in accordance with the ethical guidelines of the Institutional Animal Care and Use Committee (IACUC) of the Experimental Animal Center at Jeonbuk National University (Approval number JBNU 2026–089). Murine infection experiments were conducted in the BL3 facility of the Core Facility Center for Zoonosis Research (Core-FCZR), Jeonbuk National University. All experiments followed the experimental ethics guidelines.

### Construction of the WD40 knockout mutants

WD40 genes were deleted in the *C. neoformans* H99S strain by homologous recombination using gene disruption cassettes carrying a nourseothricin-resistant marker (nourseothricin acetyltransferase; *NAT*), generated via *NAT* split-marker/double-joint PCR strategies^42^ (Supplementary Table 2). The 5’- and 3’-flanking regions of each target gene were PCR-amplified from H99S genomic DNA using L1/L2 and R1/R2 primer pairs. The *NAT* marker, containing a signature-tagged sequence, was amplified from the pNAT-STM plasmid with M13Fe and M13Re primers. In the second round of PCR, L1/SM2 and R2/SM1 primers were used to assemble the *NAT* split-marker gene disruption cassettes. These cassettes were introduced into the H99S strain by biolistic transformation. Briefly, *C. neoformans* cells were grown in YPD medium at 30 °C for 16 h, plated on YPD agar supplemented with 1 M sorbitol, and incubated for 3 h at 30 °C. DNA-coated microcarrier beads (600 μg of 0.6-μm gold particles; Bio-Rad, Hercules, CA, USA) were loaded with the disruption cassettes and delivered into the cells using a PDS-100 particle delivery system (Bio-Rad). After a 4-h recovery at 30 °C to allow membrane repair, cells were plated on YPD agar containing nourseothricin (100 μg/ml). *NAT*-positive transformants were screened by diagnostic PCR, and genotypes were further confirmed by Southern blot analysis. At least two independent knockout mutants were obtained for each gene. The *wcp1*Δ mutants were additionally generated in *C. deneoformans* (JEC21) and *C. deuterogattii* (R265) using the same method as for *C. neoformans* H99. All primers used are listed in Supplementary Table 1.

### Construction of *WCP1* complemented, domain deletion, and epitope-tagged strains

To confirm the phenotypes of the *wcp1*Δ mutant, a *WCP1-*containing plasmid was generated using the Gibson assembly method for mutant complementation. The full-length *WCP1* gene fragment, including the 5’ flanking region (1000 bp of the native promoter), the entire ORF, and the 3’ flanking region (500 bp of the terminator), was amplified from H99 genomic DNA by Phusion PCR and cloned into the pNEO plasmid. Domain-deletion strains were generated by complementing the *wcp1*Δ mutant with truncated alleles lacking either the PPIase domain (*WCP1^PPIase^*^Δ^) or the WD40 domain (*WCP1^WD40^*^Δ^). Truncated alleles were amplified with specific primers (Supplementary Table 1) and cloned into the pNEO plasmid using Gibson assembly. For *WCP1^PPIase^*^Δ^, the PPIase-encoding region was deleted, while retaining the 5’ and 3’ flanking sequences fused to the remaining ORF. For *WCP1^WD40^*^Δ^, the WD40-encoding region was deleted in a similar manner. All constructs were linearized with MfeI and introduced into the *wcp1*Δ mutant strain (YSB10389) via biolistic transformation. Targeted re-integration of the wild-type and domain-deletion alleles at the native locus was confirmed by diagnostic PCR using specific primer sets (Supplementary Table 1). To monitor cellular localization of Wcp1 and to enable proteomics analysis, C-terminal fusions with either mCherry or a 4xFLAG epitope were constructed. The *WCP1-mCherry* and *WCP1-4xFLAG* cassettes were cloned into the pNEO plasmid and introduced into the *wcp1*Δ strain by biolistic transformation.

### Construction of *C. neoformans CTR4* promoter replacement strains

For the 37 WD40 genes that could not be deleted after multiple knockout attempts, *CTR4* promoter replacement was performed as previously described^43^. The *NAT*-*CTR4* promoter was amplified by PCR from the pNAT-CTR4 plasmid using primer pair B354/B355. Approximately 800 bp of the *CTR4* promoter region and ∼800 bp of the 5’ region of each target gene were amplified from H99 genomic DNA using primer pairs *CTR4*_L1/*CTR4_*L2 and *CTR4_*R1/*CTR4*_R2, respectively. In the first round of PCR, the promoter, 5’ exon region, and *NAT*-*CTR4* were amplified. In the second round, primer pairs *CTR4*_L1/SM2 and *CTR4*_R2/SM1 were used to generate the *NAT*-*CTR4* split-marker cassettes. The resulting insertion cassettes were introduced into *C. neoformans* H99 via biolistic transformation. Stable transformants were selected and screened for correct insertion by diagnostic PCR, and genotypes were further validated by Southern blot analysis using gene-specific probes.

### Meiotic progeny analysis for examining essentiality

To independently assess whether *WDP1* is dispensable for vegetative growth, one allele of *WDP1* was disrupted in the previously described AI187 diploid strain (*ADE2/ade2 URA5/ura5 MAT*α*/MAT***a**)^44^ using a nourseothricin-resistance marker (*NAT*). The *WDP1* disruption cassette was generated by double-joint PCR with the 5′- and 3′-flanking regions of *WDP1* and the *NAT* marker, and was introduced into the AI187 diploid strain by biolistic transformation. Stable transformants were screened by diagnostic PCR and confirmed by Southern blot analysis after BlpI digestion using a *WDP1*-specific probe. For sporulation, the heterozygous *WDP1/wdp1*Δ::*NAT* diploid strain was grown in liquid YPD medium at 30°C for 16h, washed twice with PBS and adjusted to 1 × 10^7^ cells/ml. The cell suspension was spotted onto V8 juice agar (pH 5.0) and incubated in the dark at room temperature under desiccating conditions until basidia and spore chains were visible. The sporulated cell mass—containing filaments, basidia, spores and yeast cells—was collected and suspended in 75% Percoll^®^ (Sigma Aldrich, MO, USA) made isotonic with PBS, as previously described^45^. The suspension was centrifuged at 3,000 rpm for 20min at 4°C to establish a density gradient. The spore-enriched band near the bottom of the gradient was collected, washed twice with PBS and plated onto YNB medium containing 5-fluoroorotic acid (5-FOA; 1mg/ml) to enrich for meiotic progeny and counter-select against diploid cells carrying *URA5*. Plates were incubated at room temperature for 4–5days. Recovered progeny were genotyped for mating type by PCR with mating-locus-specific primers. To assess segregation of auxotrophic and drug-resistance markers, progeny were spotted onto YPD, YPD + NAT (100µg/ml), YNB, YNB + URA (40µg/ml), YNB + ADE (20µg/ml) and YNB + URA + ADE plates. Plates were incubated at 30°C and photographed daily for 3days. Nourseothricin-resistant progeny were further analysed by internal PCR using primers annealing within the *WDP1* coding sequence.

### *Cryptococcus neoformans* WD40 Phenome Database

To enable public access to genomic and phenotypic data for WD40 domain-containing genes in *C. neoformans* H99, we developed the *Cryptococcus neoformans* WD40 Phenome Database (https://WD40.cryptococcus.org/). The database currently archives 89 WD40 genes and is implemented using MariaDB (v12.1.2) for data management and PHP (v8.4.17) for the web interface, hosted on an Apache web server in a Linux environment. Each gene entry includes detailed annotations, such as NCBI and UniProt identifiers, Pfam and KEGG orthology assignments, and external links to JBrowse^46^, FungiDB^47^, and ProteinPlus^48^, providing information on genomic localisation, functional context and protein-protein interactions. JBrowse enables contextual visualisation of gene loci together with RNA-seq expression profiles^49,50^. In addition, mutant construction designs and phenotype screening data are available for each gene. Predicted 3D structures from the AlphaFold Protein Structure Database^51^ are visualised using the NGL Viewer^52^, with WD40 domains highlighted. WD40 domains were identified using HMMER^53^ against the Pfam database (PF00400.38; canonical WD40)^54^, with a sequence-level E-value threshold < 1 and a domain-level E-value threshold < 1×10². When no canonical PF00400.38 hit was detected, subclass WD40-related Pfam domains (e.g., WD40_CDC20-Fz, WD40_WDHD1_1st and WDR55) were used for annotation.

### Growth and chemical susceptibility assays

*C. neoformans* cultures were grown in YPD medium at 30 °C for 16 h. Cells were serially diluted 10-fold (10^0^-10^4^) and spotted onto YPD agar plates containing chemical agents to induce environmental stress, as previously described^22,55,56^. Stress conditions included osmotic stress (sorbitol); cation/salt stresses (NaCl, KCl) under glucose-rich (YPD) or glucose-starved (YP) conditions; oxidative stress (H_2_O_2_, *tert*-butyl hydroperoxide, menadione, diamide); heavy metal stress (CdSO_4_); genotoxic stress (methyl methanesulfonate, hydroxyurea); membrane-destabilizing stress (sodium dodecyl sulfate); cell wall stress (calcofluor white, Congo red); ER stress (tunicamycin, dithiothreitol); and antifungal drugs (fludioxonil, fluconazole, amphotericin B, flucytosine). To assess thermotolerance and CO_2_ adaptation, serially diluted cells were spotted on YPD agar and incubated at 25 °C, 37 °C (±5% CO_2_), and 39 °C. Plates were also incubated at 30 °C for 1–5 days and photographed daily. For pH-dependent stress adaptation, YPD medium was buffered to pH 5 with 50 mM citrate (citric acid/NaOH), pH 6 with 50 mM MES, and pH 7 or 8 with 150 mM HEPES. The pH was adjusted with NaOH prior to sterilisation.

### Mating assay

To assess unilateral mating efficiency, each *MAT*α WD40 mutant (constructed in the H99S background) was crossed with the *MAT***a** KN99**a** strain. Strains were cultured in YPD medium at 30 °C for 16h, washed twice with phosphate-buffered saline (PBS), and mixed at equal concentrations (10^7^ cells/ml). Cell suspensions were spotted on V8 agar (pH 5) and incubated at 25 °C in the dark for 14 days. Filamentous growth was monitored and photographed weekly.

### In vitro virulence factor production assay

To evaluate capsule production, each WD40 mutant was cultured at 30 °C, spotted onto Dulbecco’s modified Eagle’s (DME) agar, and incubated at 37 °C for 2 days. Cells were then scraped, resuspended in distilled water, and stained with India ink (BactiDrop; Remel, San Diego, CA, USA). Capsule morphology was examined by differential interference contrast (DIC) microscopy (BX51, Olympus, Tokyo, Japan). Capsule thickness was calculated as the difference between the total cell diameter and the cell body diameter. For quantitative analysis, 50 cells were measured for each WD40 mutant and the H99S control strain. To evaluate melanin production, each WD40 mutant was grown in YPD medium at 30 °C for 16h, washed with PBS, and spotted (3μl) onto Niger seed agar containing 0.1% or 0.2% glucose. Plates were incubated at 37 °C and photographed after 1–3 days. For mutants exhibiting growth defects at 37 °C, both melanin and capsule production efficiencies were assessed at 30 °C. To evaluate urease production, each WD40 mutant was cultured at 30 °C for 16h, washed with distilled water, and inoculated (10^8^ cells) into rapid urea broth (RUH) in 10-ml medical tubes (SPL Life Sciences). Cultures were incubated at 30 °C for 2 h with shaking. Cells were pelleted by centrifugation, and the absorbance of the supernatant was measured at 570 nm using a DU 730 UV/Vis spectrophotometer (Beckman Coulter), as described previously^57^.

### STM-based murine infectivity assay

A set of WD40 mutant strains carrying unique signature-tagged NAT selection markers was first validated by PCR with tag-specific primers to confirm the presence of the corresponding barcode sequences. Each strain was cultured overnight in YPD medium at 30°C, washed three times with PBS, and adjusted to equivalent cell densities before pooling. The composition of each mutant pool is listed in Supplementary Table 3. Each pool included the virulence-positive control strain *ste50*Δ and the virulence-attenuated control strain *pka1*Δ, as previously described^58^. The pooled inoculum was prepared by combining each strain in equal proportions and diluting the mixture in PBS to a final concentration of 5 × 10^5^ cells per mouse in a total volume of 40µl. Seven-week-old female C57BL/6NCrlOri mice (Orient Bio, Seongnam, Korea) were anaesthetised by inhalation of isoflurane and intranasally infected with the pooled inoculum while spontaneous respiration was maintained in a supine position with the head tilted back, as previously described^22,55^. Five mice were used per each experimental group. At 14 days post-infection, mice were euthanised, and the lungs, brains, kidneys, livers, and spleens were harvested. Organs were homogenised in PBS, and the homogenates were plated onto YPD agar supplemented with chloramphenicol. Plates were incubated at 30°C for 3 days, and recovered fungal cells were collected by scraping. Genomic DNA was extracted from the recovered cell populations by phenol–chloroform extraction combined with bead beating. Quantitative PCR was performed with tag-specific primers (Supplementary Table 3), and STM scores were calculated by the 2^-ΔΔCt^ method to determine relative changes in strain abundance between input and output populations, as previously described^59^.

### Wcp1 murine infection model

To validate the contribution of Wcp1 and its functional domains to cryptococcal virulence using a classical murine infection model, independent infection experiments were performed using the wild-type strain, *wcp1*Δ mutant, PPIase-domain deletion strain, WD40-domain deletion strain, and the full-length *WCP1*-complemented strain. Anaesthesia, intranasal infection, and animal handling procedures were performed largely as described for the STM-based infectivity assay. Overnight cultures of *C. neoformans* strains were grown in YPD medium at 30°C, washed three times with PBS, and adjusted to a final concentration of 5×10^5^ cells in 40µl PBS per mouse. Six- to seven-week-old female BALB/cAnNCrlOri mice (Orient Bio, Seongnam, Korea) were anaesthetised with isoflurane and intranasally infected with the prepared inoculum. For survival analysis, seven mice per strain group were infected and monitored daily until predefined humane endpoints were reached. For fungal burden analysis, five mice per strain group were infected as an independent cohort and euthanised at 18 days post-infection. Lungs, brains, kidneys, livers, and spleens were harvested, homogenised in PBS, serially diluted, and plated onto YPD agar supplemented with chloramphenicol. Plates were incubated at 30°C for 3 days, and fungal burden was quantified as colony-forming units (CFU) per gram of organ tissue.

### Insect infectivity assay

*Drosophila melanogaster* (*w*^1118^) flies were used for infection experiments. Flies were maintained at 25 °C under a 12-h light/dark cycle on standard cornmeal-agar medium obtained from the Bloomington Stock Center. For infection, *Cryptococcus* strains (H99 wild-type, *wcp1*Δ mutant, complemented strain (*wcp1*Δ*::WCP1*) and domain-deletion strains (+*WCP1^PPIase^*^Δ^*, +WCP1^WD40^*^Δ^) were cultured overnight at 30 °C in YPD medium and washed once with PBS. Cell suspensions were adjusted to OD_600_ = 100 in 500 μl and supplemented with 1% (w/v) blue food dye (FD&C Blue #1) to facilitate visualisation. Adult female flies (4–5 days old) were anaesthetised with CO_2_, and the dorsal thorax was pricked with a fine needle dipped in the fungal suspension. For each strain, 50-70 flies were infected and divided into groups of 8-10 flies per vial (biological replicates). Each group was maintained in a separate vial containing fresh cornmeal-agar medium. Infected flies were incubated at 30 °C, and survival was recorded daily for up to 6 days post-infection (dpi). Kaplan–Meier survival curves were generated to compare virulence among strains.

### LysoSensor staining

Vacuolar acidification was assessed using LysoSensor Green DND-189 (Thermo Fisher Scientific, L7535). Wild-type and *wcp1*Δ strains grown overnight in YPD at 30 °C were spotted onto YPD agar plates buffered to pH 5 (50 mM citrate), 6 (50 mM MES), 7 or 8 (150 mM HEPES), and incubated for 3 days at the indicated temperature and CO_2_ conditions (30 °C, 37 °C or 37 °C with 5% CO_2_). Cells were collected from colonies, washed twice with PBS and resuspended in 1 ml of YPD containing 1 μM LysoSensor Green DND-189. Cell suspensions were incubated for 30 min in the dark with shaking (200 rpm) under the same temperature and CO_2_ conditions as the original plate. After staining, cells were pelleted, washed three times with PBS and immediately imaged by fluorescence microscopy using identical acquisition settings across all conditions. Fluorescence intensity was quantified per cell in Fiji (ImageJ) by manually outlining individual cells using corresponding DIC images. Mean fluorescence intensity per cell was measured after background subtraction. At least 50 cells per condition were quantified across three independent biological replicates and used as a relative readout of vacuolar acidification

### Macrophage intracellular survival assay

The J774A.1 murine macrophage-like cell line was obtained from the Korea Cell Line Bank (KCLB) and maintained in DMEM supplemented with 10% heat-inactivated FBS, 1% penicillin–streptomycin and 1% MEM non-essential amino acids at 37°C with 5% CO_2_. For infection assays, 1 × 10^5^ macrophages were seeded per well in 96-well plates and incubated for 16–18h to allow adherence. Macrophages were activated with 10nM phorbol 12-myristate 13-acetate (PMA) for 1h before infection. *C. neoformans* strains were grown overnight in YPD, subcultured into fresh YPD at OD_600_ = 0.3 and incubated for 16–18h at 30°C. Fungal cells were washed with PBS and adjusted to 1 × 10^6^ cells/ml. For opsonisation, 1 × 10^6^ yeast cells were incubated with 2µg anti-capsular monoclonal antibody 18B7 in macrophage medium for 1h at 37°C. Opsonised cells were added to macrophages at a multiplicity of infection (MOI) of 1. After 1h of co-culture, wells were washed three times with PBS to remove extracellular yeast, and infected macrophages were incubated in fresh medium for an additional 24h. Macrophages were then lysed with distilled water on ice for at least 30min, and lysates were serially diluted and plated onto YPD agar containing chloramphenicol (30µg/ml). CFUs were counted after 2–3days at 30°C and normalised to the verified input inoculum. Macrophage survival values were then normalised to the wild-type value set to 100%, and statistical significance was determined using a one-sample t-test against 100% in GraphPad Prism v10.4.1. Experiments were performed with three biological replicates.

### Gene expression analysis

To confirm overexpression in reciprocal genetic interaction strains, transcript levels of *KIC1*, *RIM101* and *WCP1* were measured by qRT–PCR. The *wcp1*Δ P*_H3_*:*KIC1*, *kic1*Δ P*_H3_*:*WCP1*, *wcp1*Δ P*_H3_*:*RIM101* and *rim101*Δ P*_H3_*:*WCP1* strains, together with the corresponding control strains, were cultured in liquid YPD medium at 30 °C with shaking (220 rpm) for 16 h and then subcultured into fresh YPD at an initial OD_600_ of 0.2. When cultures reached early logarithmic phase (OD_600_ = 0.6–0.8), cells were harvested by centrifugation, frozen in liquid nitrogen and lyophilised. Total RNA was extracted as previously described^60^, except that the Easy-BLUE RNA extraction kit (iNtRON Biotechnology, South Korea) was used instead of TRIzol extraction solution. cDNA was synthesised from 1 µg total RNA using RTase (Thermo Scientific, USA). qRT-PCR was performed using gene-specific primers listed in Supplementary Table 1. Transcript levels were normalised to *ACT1* and quantified using the 2^-ΔΔCt^ method, with data representing the mean of three independent biological replicates.

### Proteomics analysis of Wcp1-interacting proteins

To identify Wcp1-interactors, both Wcp1-mCherry and Wcp1-4×FLAG strains were subjected to in vivo pulldown followed by mass spectrometry-based proteomics. Strains were grown in 50 ml YPD broth at 30 °C for 16 h, subcultured into 1 L of fresh YPD, and incubated to OD_600_ 0.6-0.8. Cultures were divided into 500 ml aliquots and incubated for 1 h 30 min at pH 5 or pH 7 at 37 °C. Cells were harvested, frozen in liquid nitrogen, and lyophilised. Total proteins were extracted from lyophilised cells using lysis buffer (without SDS) containing 50 mM Tris-Cl (pH 7.5), 1% sodium deoxycholate, 5 mM sodium pyrophosphate, 0.2 mM sodium orthovanadate, 50 mM sodium fluoride, 1% Triton X-100, 0.5 mM phenylmethylsulfonyl fluoride and 2.5× protease inhibitor cocktail (Merck Millipore), following as described previously with minor modifications^61^. For the Wcp1-mCherry strain, RFP-Trap Agarose (ChromoTek, USA) was added to lysates and incubated overnight at 4 °C with rotation. For the Wcp1-4×FLAG strain, lysates were incubated with anti-FLAG antibody (Sigma-Aldrich, F1804) overnight at 4 °C, followed by incubation with Protein G Sepharose™ 4 Fast Flow (Cytiva, USA) for 6 h at 4 °C. Beads were washed three times with lysis buffer, and bound proteins were eluted in SDS sample buffer (50mM Tris-Cl, 2% SDS, 10% glycerol, 0.01% β-mercaptoethanol).

For in-gel digestion and peptide extraction, gel pieces were processed in 100 mM ammonium bicarbonate (NH_4_HCO_3_)/acetonitrile (1:1) at 2 ml per sample. Each sample was treated with 500 µl buffer and shaken for 30 min, with replacement every 30 min until fully decolourised. Samples were washed with water/acetonitrile (1:1) for 30 min and dehydrated in 100% acetonitrile until gel pieces became opaque. Samples were dried in a speed vacuum and transferred to fresh tubes. For digestion, sequencing-grade trypsin (Promega, V5111) was used at 100-200 ng per gel. Trypsin stock (100 ng/µl) was diluted to 5 ng/µl, and 20 µl was added to each sample. Samples were incubated on ice for 10 min, overlaid with 50 mM NH_4_HCO_3_ and incubated for an additional 10 min. Digestion was performed overnight at 37 °C or for 10-30 min at 37 °C using a microwave digestion system. Peptides were extracted sequentially using (i) 30 µl of 5% formic acid (FA) in 50% acetonitrile for 30-60 min, (ii) 50-100 µl of 5% FA/50% acetonitrile, and (iii) 50-100 µl of 100% acetonitrile. Supernatants were pooled, frozen, dried in a speed vacuum, and reconstituted in 20-30 µl of 0.1% FA in water.

Mass spectrometric analyses were performed using an Orbitrap Exploris 240 mass spectrometer (Thermo Scientific) coupled to a Dionex U3000 RSLCnano HPLC system. Samples were reconstituted in solvent A (water/acetonitrile, 98:2, v/v, containing 0.1% FA) and analysed by LC-nano ESI-MS/MS system. Peptides were first trapped on an Acclaim PepMap 100 trap column (100 μm x2 cm, nanoViper C18, 5 µm, 100Å, Thermo Scientific, part no. 164564) and washed with solvent A at 4 μl/min for 6 min, then separated on an IonOpticks Aurora Ultimate C18 column (25 cm × 75 μm, 1.7 μm particles, 120 Å pore size) at 300 nl/min. The LC gradient was 2–8% solvent B over 10 min, 8–30% over 55 min, followed by 90% solvent B for 4 min and re-equilibration at 2% solvent B for 20 min. Solvent B consisted of acetonitrile containing 0.1% FA. The Orbitrap analyser scanned precursor ions across 350–1,800 m/z at a resolution of 60,000 (m/z 200). Data were acquired using Xcalibur v4.4 and processed using Proteome Discoverer v2.5 (Thermo Scientific). Proteomics data have been deposited to the ProteomeXchange Consortium via the PRIDE repository^62^ under accession PXD077146 (DOI: 10.6019/PXD077146).

### Transcriptomic analysis under pH conditions

To investigate Wcp1-dependent transcriptional responses under different pH conditions, RNA-seq was performed using wild-type (H99) and *wcp1*Δ mutant strains. Cells were grown in 50 ml YPD broth at 30 °C for 16 h, subcultured into fresh YPD medium, and grown to mid-log phase (OD_600_ of 0.8). Cultures were then divided and incubated at 37 °C for 1 h 30 min under neutral (pH 7) or acidic (pH 5) conditions, matching those used for proteomic analysis. Cells were subsequently harvested, rapidly frozen in liquid nitrogen and lyophilised. Total RNA was extracted as described above. Total RNA concentration was determined using the Quant-IT RiboGreen assay (Invitrogen, #R11490). RNA integrity was assessed using the TapeStation RNA ScreenTape system (Agilent, #5067-5576), and only samples with an RNA integrity number (RIN) > 7.0 were used for library construction. Libraries were constructed independently from 1 µg total RNA per sample using the TruSeq Stranded mRNA Sample Preparation Kit (Illumina, #RS-122-2101), following the manufacturer’s instructions. Briefly, poly(A)+ mRNA was purified using oligo(dT)attached magnetic beads and fragmented using divalent cations at elevated temperature. First-strand cDNA was synthesised using SuperScript II reverse transcriptase (Invitrogen, #18064014) and random primers, followed by second-strand synthesis using DNA polymerase I, RNase H and dUTP. The resulting cDNA fragments were end-repaired, A-tailed, and ligated to sequencing adapters. Libraries were then purified and amplified by PCR. Library concentration was quantified using the KAPA Library Quantification kit for Illumina platforms (KAPA Biosystems, #KK4854), and library quality was assessed using the TapeStation D1000 ScreenTape system (Agilent, #5067-5582). Indexed libraries were sequenced on an Illumina NovaSeq X Plus platform with paired-end reads (2 × 100 bp) at Macrogen Inc.

Adapter and low-quality sequences were removed from the sequencing reads using Cutadapt (v2.4). The reference genome sequence and annotation data for *C. neoformans* H99 (GenBank assembly accession GCA_000149245.3) were obtained from the NCBI FTP server. Trimmed reads were aligned to the *C. neoformans* H99 reference genome using HISAT2 (v2.2.1). Aligned reads were converted to BAM format and sorted using Samtools (v0.1.19). Gene-level read counts were generated using featureCounts from the Subread package (v2.0.8) with the options “-T 4 -M --largestOverlap -g gene_id”. DEG analysis was performed using DESeq2 (v1.24) in R (v4.1.0). Genes with an absolute log2 fold change > 1 and a Benjamini–Hochberg-adjusted P value < 0.05 were considered differentially expressed. Volcano plots, PCA plots and heatmaps were generated using R. Alternative splicing events were analysed from HISAT2-aligned BAM files using rMATS (v4.3.0). Splice-junction usage was quantified based on inclusion junction counts, skipping junction counts, inclusion levels and inclusion-level differences. Events with an FDR < 0.05 were considered significant. Representative splice-junction changes were visualised as sashimi plots. RNA-seq data generated in this study have been deposited in the Gene Expression Omnibus under accession number GSE328123.

## Supporting information

Inventory of Supporting Information

## Acknowledgements

This work was supported by the National Research Foundation of Korea (NRF) grant funded by the Korean government (MSIT) (RS-2025-18362970, RS-2025-00555365, and RS-2025-02215093 to YSB; RS-2022-NR072215 and RS-2021-NF000550 to KTL; RS-2024-00345184 to WJL). This research was also partly supported by the Strategic Initiative for Microbiomes in Agriculture and Food funded by Ministry of Agriculture, Food and Rural Affairs (RS-2018-IP918012 to Y.-S.B.). This work was supported in part by AmtixBio Co., Ltd. (to Y.-S.B.). We thank Dr. Xiaorong Lin and Dr. Damian J. Krysan for helpful discussions. The funders had no role in study design, data collection and analysis, decision to publish, or preparation of the manuscript. BioRender was used to create schematics in Figs. 1a, 2a,b, 4a, 5a,b and 6e and Extended Data Fig. 2a; publication license: Choi, J. (2026) https://BioRender.com/zhql9du

## Data availability

All data supporting the findings of this study are available within the Article, Extended Data, Supplementary Information and Source Data files. Gene annotation, mutant construction and phenotypic profiling data are available through the *C. neoformans* WD40 Phenome Database at https://WD40.cryptococcus.org/. RNA-seq data have been deposited in GEO under accession GSE328123. Proteomics data have been deposited in PRIDE/ProteomeXchange under accession PXD077146 and DOI 10.6019/PXD077146. Source data not covered by the WD40 Phenome Database have been deposited in figshare at https://doi.org/10.6084/m9.figshare.32253381. All deposited datasets will be publicly available upon publication.

## Author contributions

Y.-S.B. and K.-T.L. conceived the project. J.-T.C., S.-R.Y., J.O., Y.-B.J., Y.L., H.C., D.W., D.K., S.Y., S.Y., E.-S.K., S.K., C.K., and K.-A.L. performed experiments and analysed the data. J.-S.L., J.C., W.-J.L., K.-T.L. and Y.-S.B. provided advice on the project and supervised the experimental analysis. J.-T.C., K.-T.L. and Y.-S.B. wrote the manuscript. All authors reviewed and approved this manuscript.

## References

1 Stelzl, U. et al. A human protein-protein interaction network: a resource for annotating the proteome. Cell 122, 957–968 (2005).

2 Mangani, S. Disruption of protein-protein interfaces: in search of new inhibitors. (Springer, 2013).

3 Rual, J. F. et al. Towards a proteome-scale map of the human protein-protein interaction network. Nature 437, 1173–1178 (2005).

4 Nicod, C., Banaei-Esfahani, A. & Collins, B. C. Elucidation of host-pathogen protein-protein interactions to uncover mechanisms of host cell rewiring. Curr Opin Microbiol 39, 7–15 (2017).

5 Lu, H. et al. Recent advances in the development of protein-protein interactions modulators: mechanisms and clinical trials. Signal Transduct Target Ther 5, 213 (2020).

6 Kuntz, I. D. Structure-based strategies for drug design and discovery. Science 257, 1078–1082 (1992).

7 Hopkins, A. L. & Groom, C. R. The druggable genome. Nat Rev Drug Discov 1, 727–730 (2002).

8 Congreve, M., Murray, C. W. & Blundell, T. L. Structural biology and drug discovery. Drug Discov Today 10, 895–907 (2005).

9 Stirnimann, C. U., Petsalaki, E., Russell, R. B. & Muller, C. W. WD40 proteins propel cellular networks. Trends Biochem Sci 35, 565–574 (2010).

10 Jain, B. P. & Pandey, S. WD40 Repeat Proteins: Signalling Scaffold with Diverse Functions. Protein J 37, 391–406 (2018).

11 Schapira, M., Tyers, M., Torrent, M. & Arrowsmith, C. H. WD40 repeat domain proteins: a novel target class? Nat Rev Drug Discov 16, 773–786 (2017).

12 Bloom, A. L. M. et al. Thermotolerance in the pathogen *Cryptococcus neoformans* is linked to antigen masking via mRNA decay-dependent reprogramming. Nat Commun 10, 4950 (2019).

13 Chadwick, B. J., Ristow, L. C., Xie, X., Krysan, D. J. & Lin, X. Discovery of CO(2) tolerance genes associated with virulence in the fungal pathogen *Cryptococcus neoformans*. Nat Microbiol 9, 2684–2695 (2024).

14 Dragotakes, Q. et al. Macrophages use a bet-hedging strategy for antimicrobial activity in phagolysosomal acidification. J Clin Invest 130, 3805–3819 (2020).

15 Alvarez, M. & Casadevall, A. Phagosome extrusion and host-cell survival after *Cryptococcus neoformans* phagocytosis by macrophages. Curr Biol 16, 2161–2165 (2006).

16 Denning, D. W. Global incidence and mortality of severe fungal disease. Lancet Infect Dis 24, e428–e438 (2024).

17 Ma, J. et al. WDSPdb: an updated resource for WD40 proteins. Bioinformatics 35, 4824–4826 (2019).

18 Zou, X. D. et al. Genome-wide Analysis of WD40 Protein Family in Human. Sci Rep 6, 39262 (2016).

19 Yang, D. H. et al. Pleiotropic roles of the Msi1-like protein Msl1 in *Cryptococcus neoformans*. Eukaryot Cell 11, 1482–1495 (2012).

20 Liu, R., et al. The COMPASS Complex Regulates Fungal Development and Virulence through Histone Crosstalk in the Fungal Pathogen Cryptococcus neoformans. J Fungi (Basel) 9 (2023).

21 Peterson, P. P. et al. The *Cryptococcus neoformans* STRIPAK complex controls genome stability, sexual development, and virulence. PLoS Pathog 20, e1012735 (2024).

22 Lee, K. T. et al. Systematic functional analysis of kinases in the fungal pathogen *Cryptococcus neoformans*. Nat Commun 7, 12766 (2016).

23 Dumesic, P. A. et al. Product binding enforces the genomic specificity of a yeast polycomb repressive complex. Cell 160, 204–218 (2015).

24 Wang, P., Perfect, J. R. & Heitman, J. The G-protein beta subunit GPB1 is required for mating and haploid fruiting in *Cryptococcus neoformans*. Mol Cell Biol 20, 352–362 (2000).

25 Ero, R. et al. Crystal structure of Gib2, a signal-transducing protein scaffold associated with ribosomes in *Cryptococcus neoformans*. Sci Rep 5, 8688 (2015).

26 Lee, H., Chang, Y. C., Varma, A. & Kwon-Chung, K. J. Regulatory diversity of *TUP1* in *Cryptococcus neoformans*. Eukaryot Cell 8, 1901–1908 (2009).

27 Wu, T., Fan, C. L., Han, L. T., Guo, Y. B. & Liu, T. B. Role of F-box Protein Cdc4 in Fungal Virulence and Sexual Reproduction of *Cryptococcus neoformans*. Front Cell Infect Microbiol 11, 806465 (2021).

28 Ory, J. J., Griffith, C. L. & Doering, T. L. An efficiently regulated promoter system for *Cryptococcus neoformans* utilizing the CTR4 promoter. Yeast 21, 919–926 (2004).

29 Chadwick, B. J. et al. The RAM signaling pathway links morphology, thermotolerance, and CO(2) tolerance in the global fungal pathogen *Cryptococcus neoformans*. Elife 11 (2022).

30 Henry, R. P. & Swenson, E. R. The distribution and physiological significance of carbonic anhydrase in vertebrate gas exchange organs. Respir Physiol 121, 1–12 (2000).

31 Geers, C. & Gros, G. Carbon dioxide transport and carbonic anhydrase in blood and muscle. Physiol Rev 80, 681–715 (2000).

32 Weitzel, G., Pilatus, U. & Rensing, L. The cytoplasmic pH, ATP content and total protein synthesis rate during heat-shock protein inducing treatments in yeast. Exp Cell Res 170, 64–79 (1987).

33 Triandafillou, C. G., Katanski, C. D., Dinner, A. R. & Drummond, D. A. Transient intracellular acidification regulates the core transcriptional heat shock response. Elife 9 (2020).

34 Ost, K. S., O’Meara, T. R., Huda, N., Esher, S. K. & Alspaugh, J. A. The *Cryptococcus neoformans* alkaline response pathway: identification of a novel rim pathway activator. PLoS Genet 11, e1005159 (2015).

35 Billmyre, R. B. et al. Landscape of essential growth and fluconazole-resistance genes in the human fungal pathogen *Cryptococcus neoformans*. PLoS Biol 23, e3003184 (2025).

36 Padrick, S. B., Doolittle, L. K., Brautigam, C. A., King, D. S. & Rosen, M. K. Arp2/3 complex is bound and activated by two WASP proteins. Proc Natl Acad Sci U S A 108, E472–479 (2011).

37 Hu, G. et al. PI3K signaling of autophagy is required for starvation tolerance and virulenceof *Cryptococcus neoformans*. J Clin Invest 118, 1186–1197 (2008).

38 Bahn, Y. S., Cox, G. M., Perfect, J. R. & Heitman, J. Carbonic anhydrase and CO_2_ sensing during *Cryptococcus neoformans* growth, differentiation, and virulence. Curr Biol 15, 2013–2020 (2005).

39 Aranda-Chan, V. et al. Insights into Peptidyl-Prolyl cis-trans Isomerases from Clinically Important Protozoans: From Structure to Potential Biotechnological Applications. Pathogens 13 (2024).

40 Rajan, S. & Yoon, H. S. Structural insights into Plasmodium PPIases. Front Cell Infect Microbiol 12, 931635 (2022).

41 Pemberton, T. J. Identification and comparative analysis of sixteen fungal peptidyl-prolyl cis/trans isomerase repertoires. BMC Genomics 7, 244 (2006).

42 Jung, K. W., Lee, K. T., So, Y. S. & Bahn, Y. S. Genetic manipulation of *Cryptococcus neoformans*. Curr Protoc Microbiol 50, e59 (2018).

43 Kim, M. S., Kim, S. Y., Jung, K. W. & Bahn, Y. S. Targeted gene disruption in *Cryptococcus neoformans* using double-joint PCR with split dominant selectable markers. Methods Mol Biol 845, 67–84 (2012).

44 Ianiri, G. & Idnurm, A. Essential gene discovery in the basidiomycete Cryptococcus neoformans for antifungal drug target prioritization. mBio 6 (2015).

45 Botts, M. R., Giles, S. S., Gates, M. A., Kozel, T. R. & Hull, C. M. Isolation and characterization of *Cryptococcus neoformans* spores reveal a critical role for capsule biosynthesis genes in spore biogenesis. Eukaryot Cell 8, 595–605 (2009).

46 Diesh, C. et al. JBrowse 2: a modular genome browser with views of synteny and structural variation. Genome biology 24, 74 (2023).

47 Basenko, E. Y. et al. What is new in FungiDB: a web-based bioinformatics platform for omics-scale data analysis for fungal and oomycete species. Genetics 227, iyae035 (2024).

48 Ehrt, C. et al. Proteins Plus: a publicly available resource for protein structure mining. Nucleic Acids Research 53, W478–W484 (2025).

49 Jang, E.-H., Kim, J.-S., Yu, S.-R. & Bahn, Y.-S. Unraveling capsule biosynthesis and signaling networks in *Cryptococcus neoformans*. Microbiology spectrum 10 (2022).

50 Do, E., Cho, Y.-J., Kim, D., Kronstad, J. W. & Jung, W. H. A transcriptional regulatory map of iron homeostasis reveals a new control circuit for capsule formation in *Cryptococcus neoformans*. Genetics 215, 1171–1189 (2020).

51 Varadi, M. et al. AlphaFold Protein Structure Database in 2024: providing structure coverage for over 214 million protein sequences. Nucleic acids research 52, D368–D375 (2024).

52 Rose Alexander, S., Bradley Anthony, R., Valasatava Yana, D. J. M. & Prlić Andreas, R. P. W. NGL viewer: web-based molecular graphics for large complexes Bioinformatics. Volume 34, 3755–3758 (2018).

53 Eddy, S. R. Accelerated profile HMM searches. PLoS computational biology 7, e1002195 (2011).

54 Mistry, J. et al. Pfam: The protein families database in 2021. Nucleic acids research 49, D412–D419 (2021).

55 Jung, K. W. et al. Systematic functional profiling of transcription factor networks in *Cryptococcus neoformans*. Nat Commun 6, 6757 (2015).

56 Jin, J. H. et al. Genome-wide functional analysis of phosphatases in the pathogenic fungus *Cryptococcus neoformans*. Nat Commun 11, 4212 (2020).

57 Kwon-Chung, K. J., Wickes, B. L., Booth, J. L., Vishniac, H. S. & Bennett, J. E. Urease inhibition by EDTA in the two varieties of *Cryptococcus neoformans*. Infect Immun 55, 1751–1754 (1987).

58 Jung, K. W., Lee, K. T. & Bahn, Y. S. A Signature-Tagged Mutagenesis (STM)-based murine-infectivity assay for *Cryptococcus neoformans*. J Microbiol 58, 823–831 (2020).

59 Livak, K. J. & Schmittgen, T. D. Analysis of relative gene expression data using real-time quantitative PCR and the 2(-Delta Delta C(T)) Method. Methods 25, 402–408 (2001).

60 Ko, Y. J. et al. Remodeling of global transcription patterns of *Cryptococcus neoformans* genes mediated by the stress-activated HOG signaling pathways. Eukaryot Cell 8, 1197–1217 (2009).

61 Choi, J. T. et al. The hybrid RAVE complex plays V-ATPase-dependent and - independent pathobiological roles in *Cryptococcus neoformans*. PLoS Pathog 19, e1011721 (2023).

62 Perez-Riverol, Y. et al. The PRIDE database at 20 years: 2025 update. Nucleic Acids Res 53, D543–D553 (2025).

