## Supplementary material for "Systematic profiling of WD40 proteins reveals Wcp1, a cyclophilin linking CO_2_/heat tolerance to acidic pH adaptation in *Cryptococcus neoformans*": Inventory of Supporting Information

**Inventory of Supporting Information for**

**This PDF file includes:**

**Extended Data Fig. 1–7 legends**

**Supplementary Table 1–3 description**

**Supplementary Data 1–7 description**

### Extended Data Figure Legends

**Extended Data Fig. 1: Evolutionary conservation of canonical WD40 proteins and antifungal hypersensitivity of selected *CTR4* promoter-replacement strains. a,** Comparative BLAST matrix analysis of canonical WD40 genes using the Comparative Fungal Genome Platform (http://cfgp.riceblast.snu.ac.kr). **Abbreviations**: Pi, *Phytophthora infestans*; Af, *Aspergillus fumigatus*; An, *Aspergillus nidulans*; Bg, *Blumeria graminis*; Bc, *Botrytis cinerea*; Ci, *Coccidioides immitis*; Cg, *Colletotrichum graminicola*; Fg, *Fusarium graminearum*; Fo, *Fusarium oxysporum*; Hc, *Histoplasma capsulatum*; Mo, *Magnaporthe oryzae*; Mg, *Mycosphaerella graminicola*; Nc, *Neurospora crassa*; Pa, *Podospora anserina*; Ca, *Candida albicans*; Sc, *Saccharomyces cerevisiae*; Sp, *Schizosaccharomyces pombe;* Cn, *Cryptococcus neoformans*; Hi, *Heterobasidion irregulare*; Lb, *Laccaria bicolour*; Pc, *Phanerochaete chrysosporium*; Sl, *Serpula lacrymans*; Ml, *Melampsora laricis-populina*; Pg, *Puccinia graminis*; Um, *Ustilago maydis*; Am, *Allomyces macrogynus*; Bd, *Batrachochytrium dendrobatidis*; Ec, *Encephalitozoon cuniculi*; Pb, *Phycomyces blakesleeanus*; Ro, *Rhizopus oryzae*; Dm, *Drosophila melanogaster*; Hs, *Homo sapiens*; Ce, *Caenorhabditis elegans*; At, *Arabidopsis thaliana*; Os, *Oryza sativa*. Light blue font indicates putative essential WD40 gene. **b,** Serial-dilution growth analysis of selected *CTR4* promoter-replacement strains under basal or copper-repressive conditions in the absence or presence of sublethal concentrations of antifungal agents: amphotericin B (AMB, 0.8 μg/ml), fluconazole (FCZ, 10 μg/ml), flucytosine (5FC, 300 μg/ml) and fludioxonil (FDX, 1 μg/ml).

**Extended Data Fig. 2: AI187-based meiotic progeny analysis confirms that *WDP1* is dispensable for growth. a**, Schematic overview of the AI187-based meiotic progeny analysis workflow used to assess *WDP1* essentiality. A heterozygous *WDP1/wdp1*Δ*::NAT* diploid strain was generated, induced to sporulate on V8 medium and subjected to Percoll-based spore enrichment and meiotic progeny analysis. **b,** Strategy and validation for construction of the heterozygous *WDP1/wdp1*Δ*::NAT* diploid strain. One allele of *WDP1* was replaced with the nourseothricin-resistance cassette in the AI187 background. The knockout design, restriction sites and probe positions used for Southern blot analysis are indicated. Southern blot analysis after BlpI digestion confirmed disruption of one *WDP1* allele while retaining the wild-type allele. **c,** Microscopic observation of spore chains from the heterozygous *WDP1/wdp1*Δ*::NAT* diploid strain. Scale bar, 20 µm. **d,** Meiotic progeny screening. Spores were enriched by Percoll gradient centrifugation and selected on 5-FOA-containing YNB medium. Recovered progeny were tested for mating type, auxotrophic marker segregation, nourseothricin resistance and growth on the indicated media. Plates were incubated at 30 °C and photographed daily for 3 days. **e,** Internal PCR verification of nourseothricin-resistant progeny. Among 21 nourseothricin-resistant progeny analysed, 3 retained an internal *WDP1* PCR product, whereas 18 lacked the internal PCR product. A minus symbol indicates loss of the internal *WDP1* region, confirming successful knockout of the *WDP1* allele. Recovery of viable nourseothricin-resistant, internal PCR-negative progeny indicates that *WDP1* is non-essential.

**Extended Data Fig. 3: Construction and validation of *WCP1* complementation, domain-deletion, epitope-tagged and reciprocal overexpression strains. a,** Schematic representation and diagnostic PCR validation of *WCP1* complementation strains constructed in the *wcp1*Δ::*NAT* background. The native *WCP1* locus, the *wcp1*Δ::*NAT* mutant allele, and the re-integrated *WCP1* alleles are shown, including full-length *WCP1*, $WCP1^{PPIase\Delta}$, $WCP1^{WD40\Delta}$, *WCP1-mCherry* and *WCP1-4×FLAG*. Primer positions used for diagnostic PCR are indicated in the schematic. Targeted integration of each construct was confirmed by diagnostic PCR using the indicated primer sets. Expected PCR product sizes are shown below each gel image.

**b**, Wcp1-mCherry and Wcp1-4×FLAG strains exhibited recovery identical to that of the wild-type strain. Plates were photographed on day 3. **c,** Construction and validation of reciprocal overexpression strains used to assess the genetic interaction between *WCP1* and *KIC1*. The $P_{H3}$:*KIC1* construct was introduced into the *wcp1*Δ background, and the $P_{H3}$:*WCP1* construct was introduced into the *kic1*Δ background. Correct integration was verified by Southern blot analysis, and overexpression of *KIC1* or *WCP1* was confirmed by qRT–PCR.

**Extended Data Fig. 4: pH-dependent growth defect and intracellular acidification phenotype of *wcp1*Δ. a,** Spot assays of WT, *wcp1*Δ, *+WCP1*, *+WCP1*$^{PPIase\Delta}$, *+WCP1*$^{WD40\Delta}$, *+WCP1-mCherry* and *+WCP1-4×FLAG* strains on unbuffered or pH-adjusted RPMI under 30 °C, 37 °C and 37 °C + 5% $CO_2$ conditions. **b**, Representative CFU assay plates of WT and *wcp1*Δ at pH 5 and pH 7 under the indicated temperature and $CO_2$ conditions. **c**, Quantification of LysoSensor fluorescence intensity in WT and *wcp1*Δ across the indicated pH, temperature and $CO_2$ conditions. Data are combined from three biologically independent experiments.

**Extended Data Fig. 5: Genetic interaction analysis between *WCP1* and *RIM101*. a,** Design and validation of the strains used for reciprocal overexpression and double-mutant analysis, including *wcp1*Δ $P_{H3}$:*RIM101*, *rim101*Δ $P_{H3}$:*WCP1*, and *wcp1*Δ *rim101*Δ. Correct integration of the $P_{H3}$:*RIM101*, $P_{H3}$:*WCP1* and *rim101*Δ::HYG alleles was verified by Southern blot analysis using the indicated restriction enzymes and probes. Overexpression of *RIM101* in the *wcp1*Δ background and *WCP1* in the *rim101*Δ background was confirmed by qRT–PCR. **b**, Spot assays of WT, *wcp1*Δ, *wcp1*Δ $P_{H3}$:*RIM101*, *rim101*Δ, *rim101*Δ $P_{H3}$:*WCP1*, and *wcp1*Δ *rim101*Δ strains on unbuffered or pH-adjusted YPD under 30 °C, 37 °C and 37 °C + 5% $CO_2$ conditions. Plates were photographed on day 4.

**Extended Data Fig. 6: Stress- and carbon-source-dependent phenotypes of *wcp1*Δ. a,** Stress- and carbon-source-dependent growth phenotyping of WT, *wcp1*Δ and +*WCP1* strains under the indicated temperature, $CO_2$, pH and medium conditions. Plates were photographed on day 4. **b**, Spot assays of WT, *wcp1*Δ and +*WCP1* strains on YNB containing 2% glucose or 1% acetic acid at pH 5 or pH 7 under 30 °C, 37 °C and 37 °C + 5% $CO_2$ conditions. YNB + glucose plates were photographed on day 4, and YNB + acetic acid plates were photographed on day 10.

**Extended Data Fig. 7: Transcriptomic and splicing features of the Wcp1-dependent acidification response. a,** Principal component analysis of transcriptomes from WT and *wcp1*Δ cells under neutral and acidic conditions. **b**, GO enrichment analysis of differentially expressed genes across the indicated transcriptomic comparisons. **c**, Functional annotation summary of significantly altered genes in each comparison, including annotated genes, hypothetical proteins and unspecified products. **d**, Numbers of significantly altered alternative splicing events in the indicated genotype and pH comparisons. **e**, Representative sashimi plot showing a Wcp1-dependent alternative splicing event in CNAG_01295 under acidic conditions.

**Supplementary Table description**

**Supplementary Table 1**
Description: List of primers used in this study, including general primers, primers for WD40 knockout mutant construction and primers for CTR4 promoter-replacement strain construction.

**Supplementary Table 2**
Description: List of *Cryptococcus neoformans* strains used in this study, including WD40 knockout mutants and putative essential WD40 CTR4 promoter-replacement strains.

**Supplementary Table 3**
Description: STM mutant pools and signature-tag-specific primers used for STM-based murine infectivity assays.

**Supplementary Data description**

**Supplementary Data 1**
Description: List and annotation of 140 putative WD40 genes identified in *Cryptococcus neoformans*, including gene identifiers, orthologues, protein sequences, InterPro annotations and confidence categories.

**Supplementary Data 2**
Description: Lists of putative WD40 proteins in representative fungal species, including *Schizosaccharomyces pombe*, *Saccharomyces cerevisiae*, *Candida albicans* and *Cryptococcus neoformans*, with repeat number, confidence category and essentiality information.

**Supplementary Data 3**
Description: Comparative BLAST and E-value matrix analysis of putative WD40 proteins in *Cryptococcus neoformans* and other eukaryotic species.

**Supplementary Data 4**
Description: Complete in vitro phenome heatmap dataset for *C. neoformans* WD40 mutants across growth, virulence-factor, stress-response and antifungal-susceptibility conditions.

**Supplementary Data 5**
Description: Pathogenicity-related WD40 proteins identified in *C. neoformans*, including functional annotations and organ-specific STM infectivity scores.

**Supplementary Data 6**
Description: Wcp1-associated candidate proteins identified by AP–MS, grouped by pH specificity as pH 5-specific, pH 7-specific or shared interactors.

**Supplementary Data 7**
Description: Differentially expressed genes identified by RNA-seq analysis under pH 5 and pH 7 conditions, with replicate-level read counts, statistical values and integrated Wcp1 pulldown overlap annotations.
