## Supplementary Figures for "Systematic profiling of WD40 proteins reveals Wcp1, a cyclophilin linking CO_2_/heat tolerance to acidic pH adaptation in *Cryptococcus neoformans*"

#### Extended Data Figure 1 (Choi et. al.)

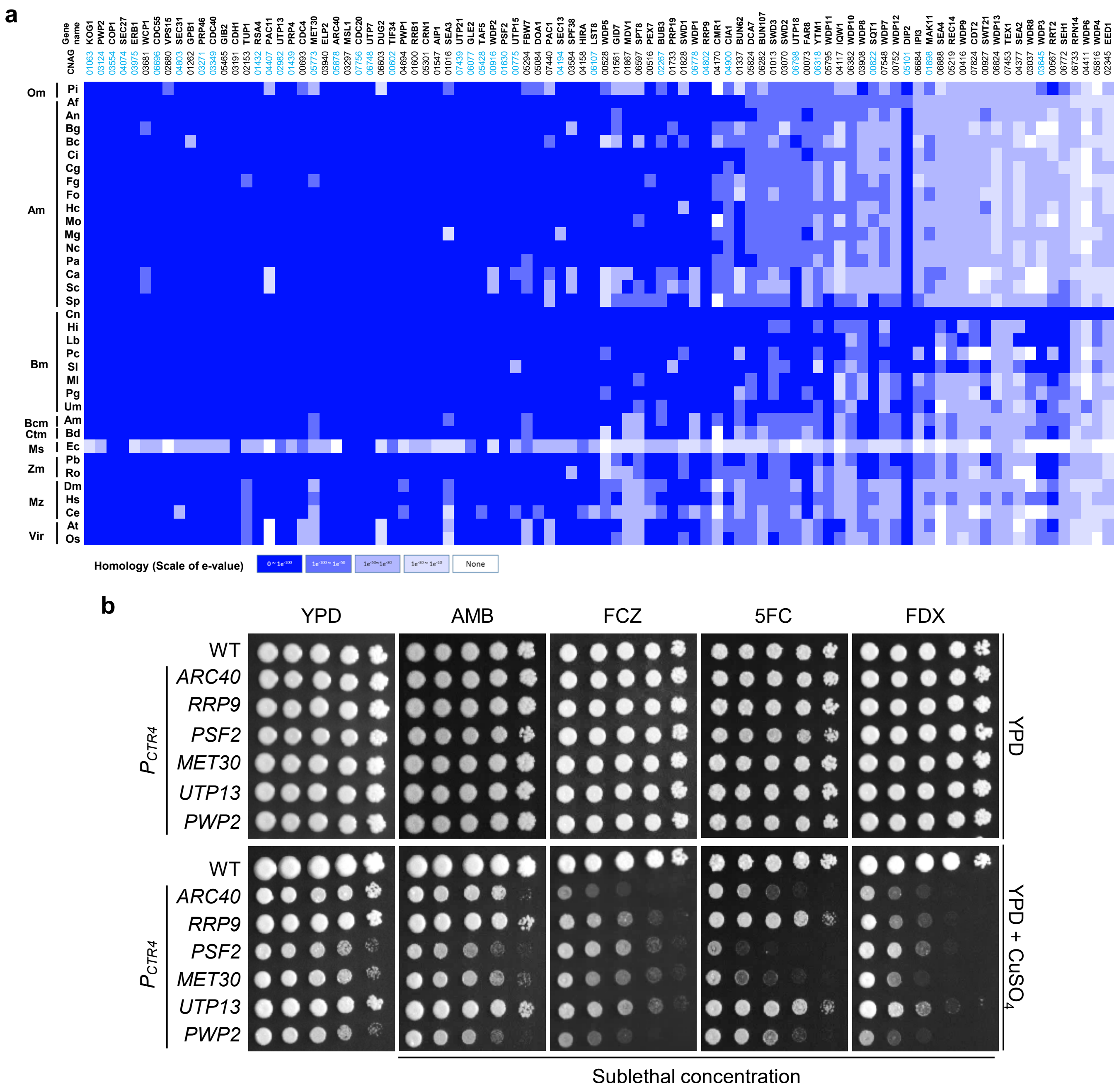

#### Extended Data Figure 2 (Choi et. al.)

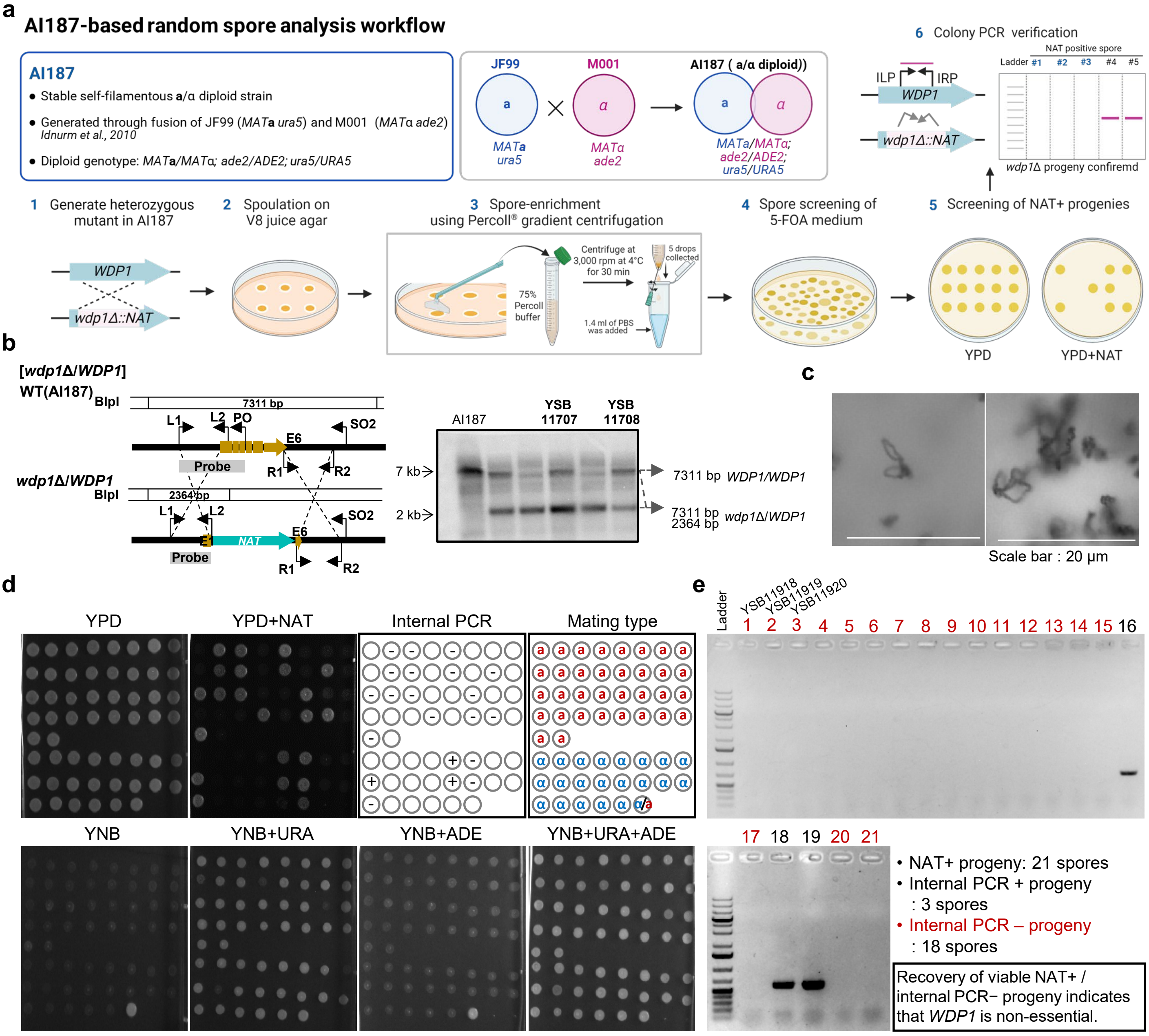

**Extended Data Figure 3 (Choi et. al.)**

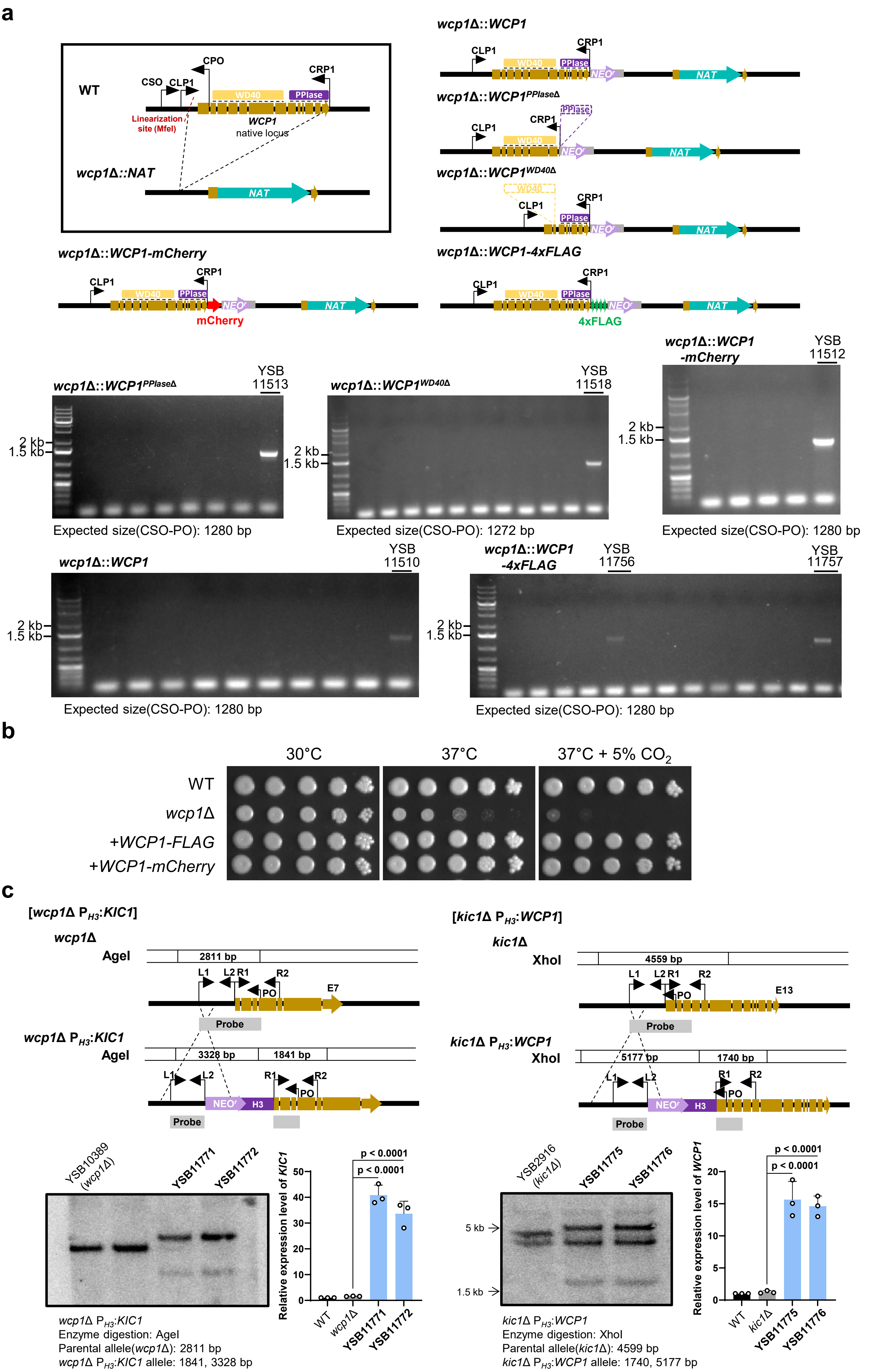

### Extended Data Figure 4 (Choi et. al.)

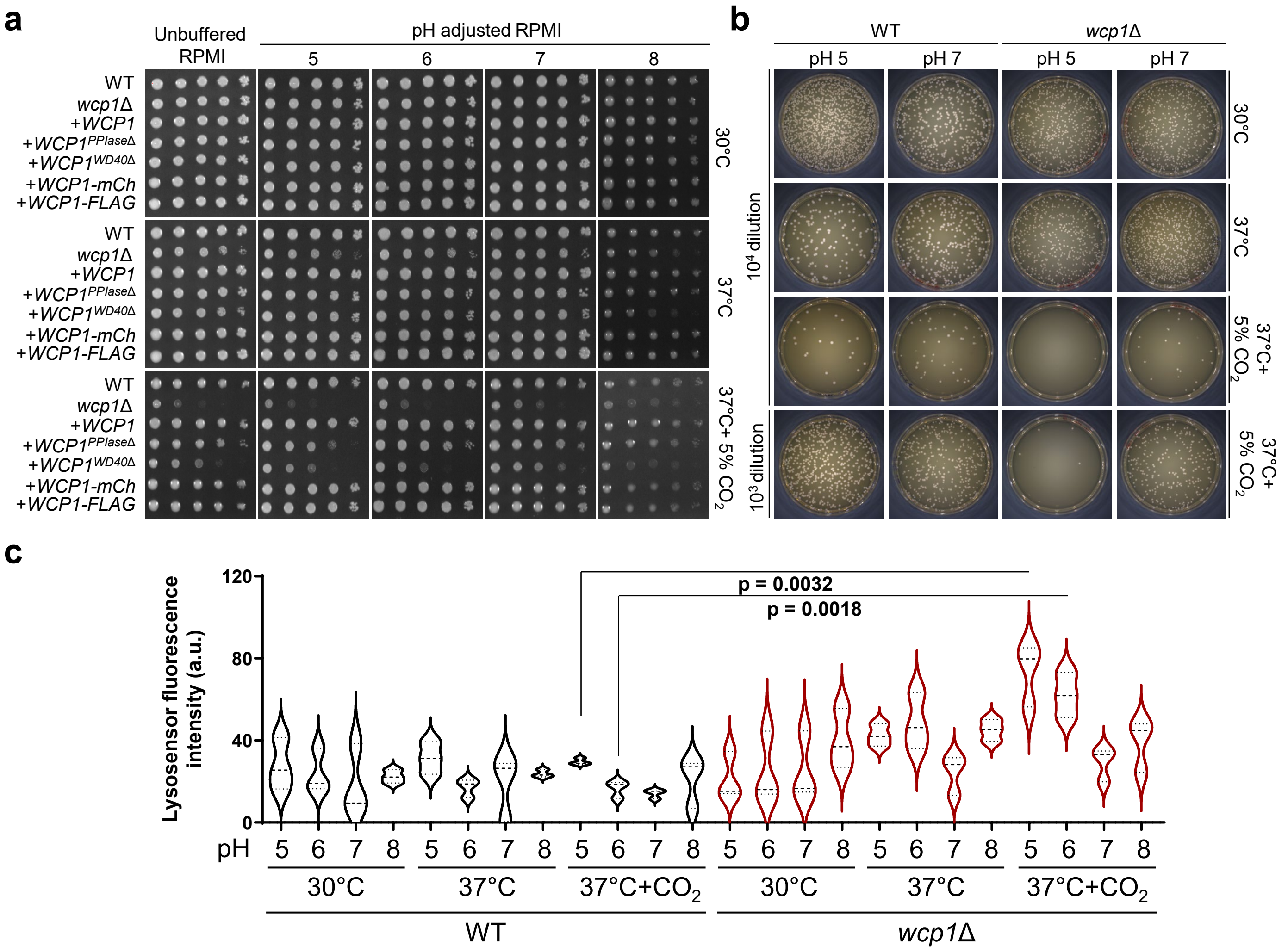

#### Extended Data Figure 5 (Choi et. al.)

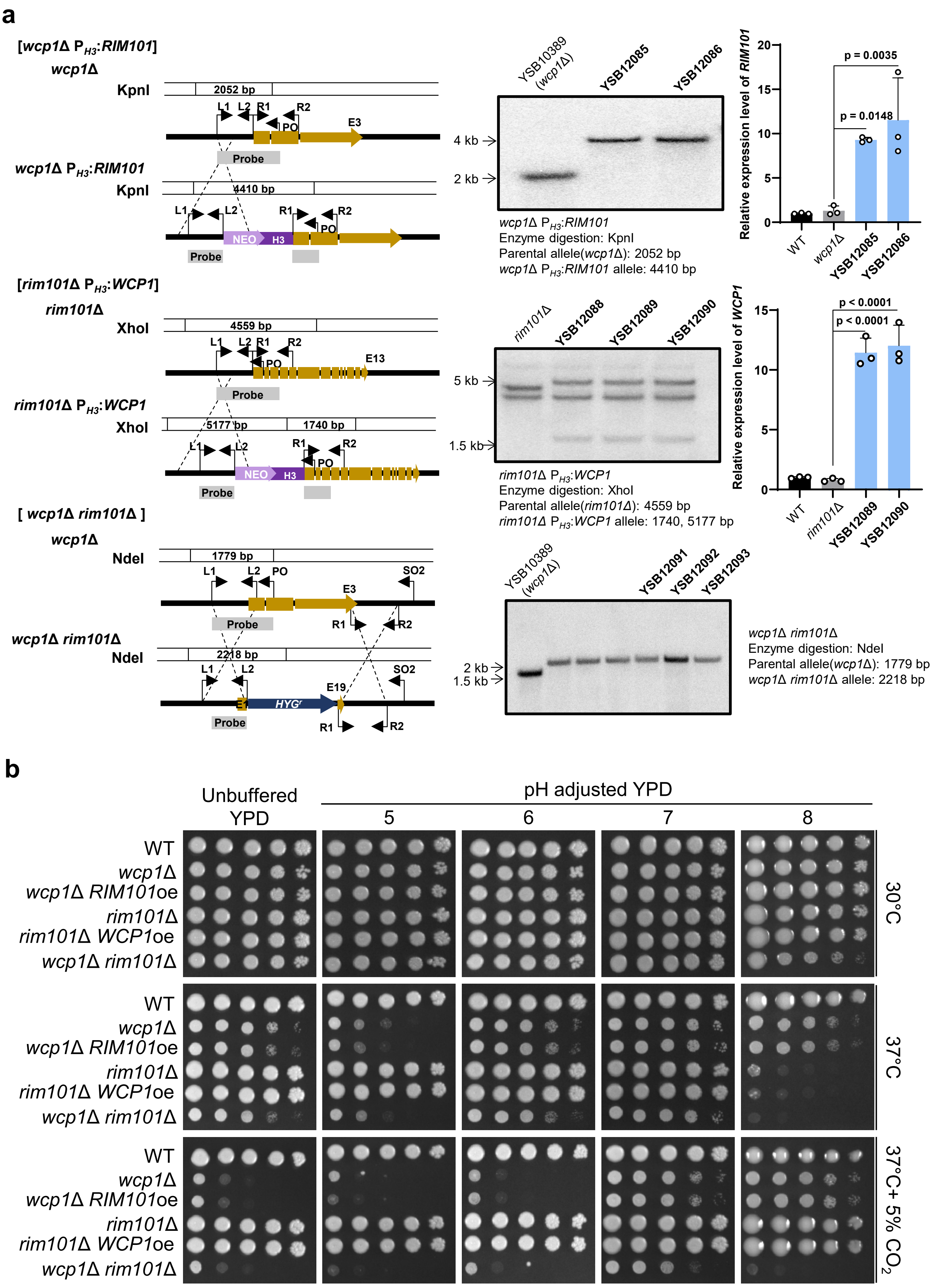

**Extended Data Figure 6 (Choi et. al.)**

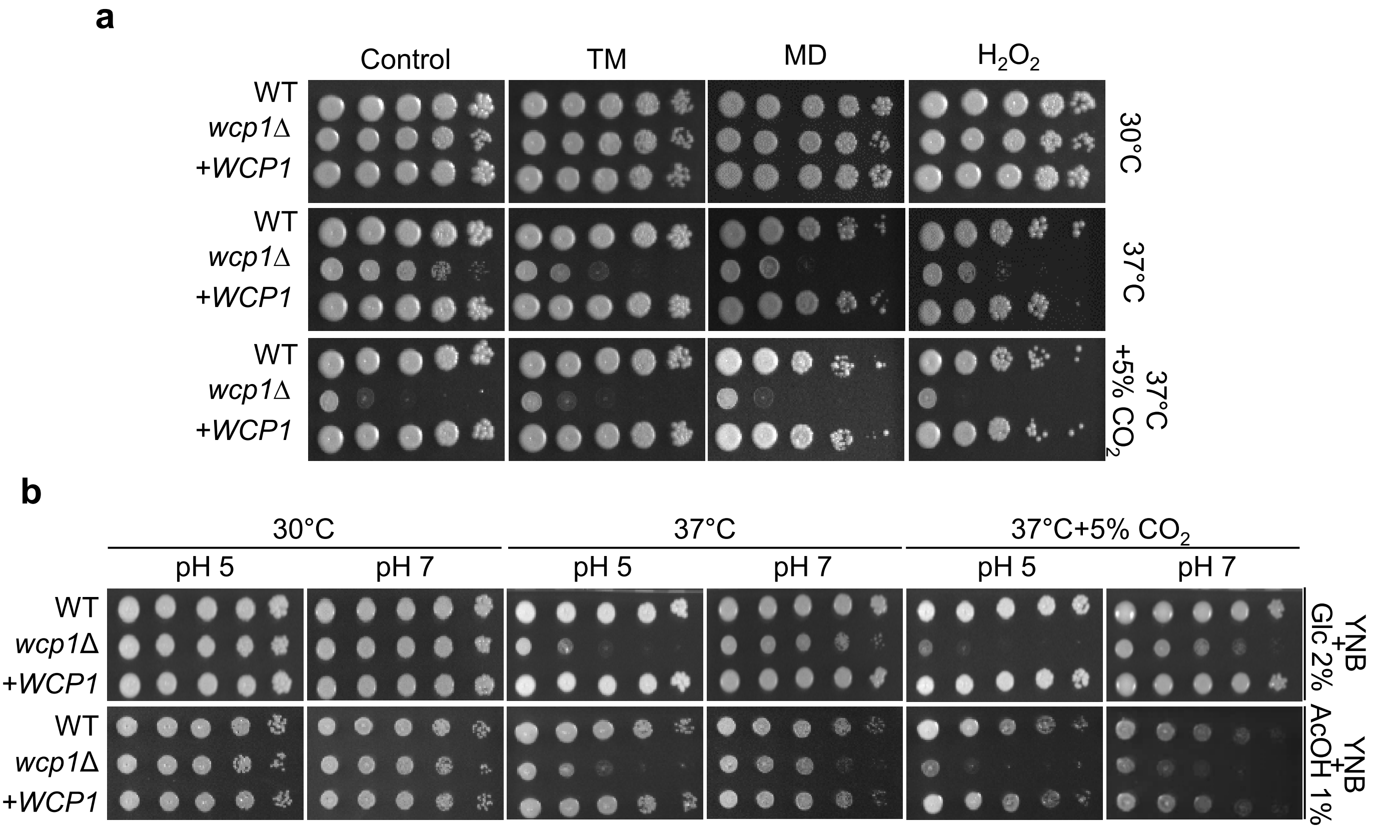

### Extended Data Figure 7 (Choi et. al.)

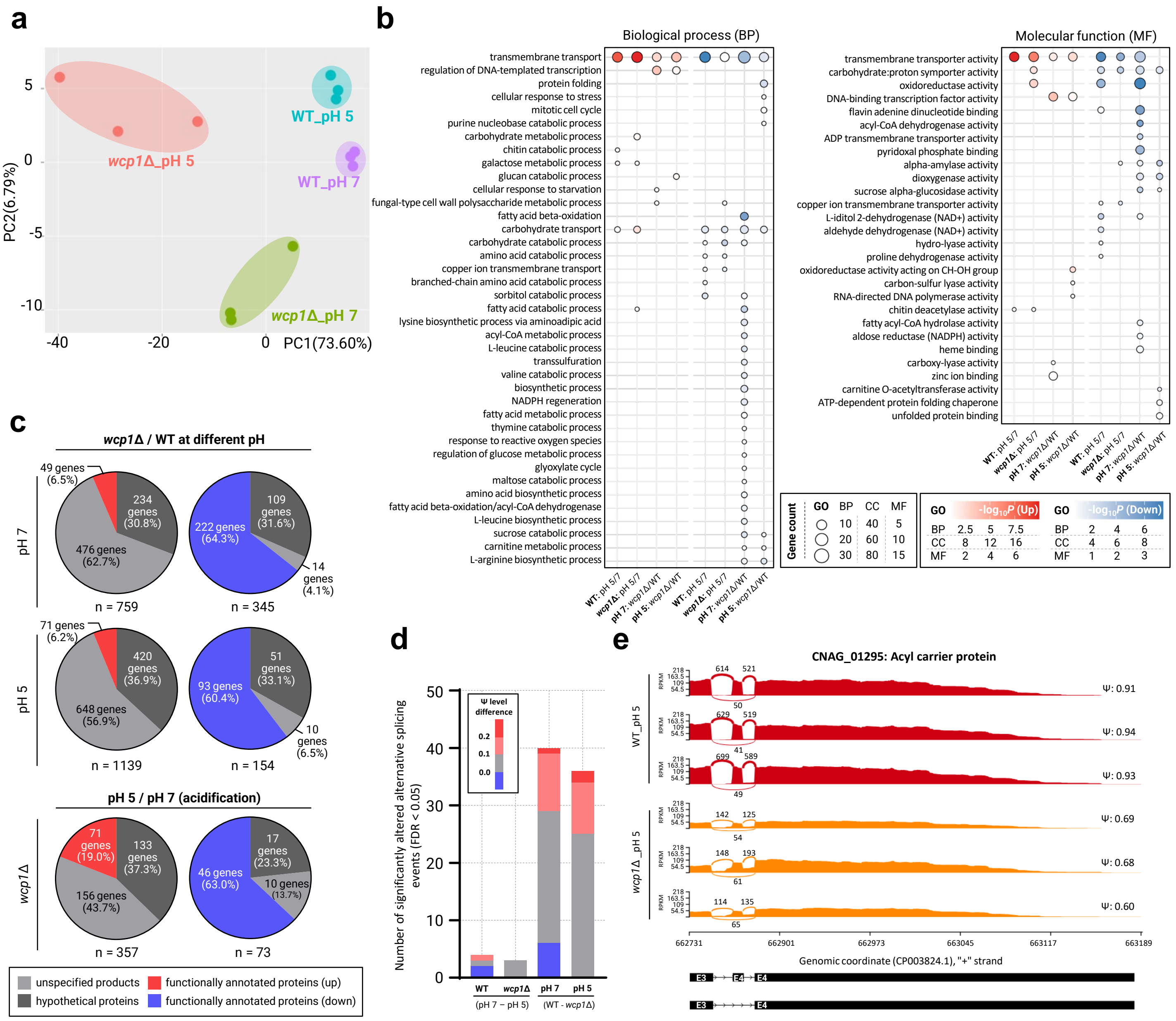
